# Determination of Cas9/dCas9 associated toxicity in microbes

**DOI:** 10.1101/848135

**Authors:** Chitra Seetharam Misra, Gargi Bindal, Megha Sodani, Surbhi Wadhawan, Savita Kulkarni, Satyendra Gautam, Rita Mukhopadhyaya, Devashish Rath

## Abstract

The CRISPR-Cas9 system has been used extensively in eukaryotic and prokaryotic systems for various applications. In case of the latter, a couple of previous studies had shown Cas9 protein expression associated toxicity. We studied the same in five microbes, viz *Escherichia coli, Salmonella typhimurium, Mycobacterium smegmatis, Xanthomonas campestris* and *Deinococcus radiodurans*. Transformation efficiency of plasmids carrying genes coding for Cas9 or dCas9 was used to gauge toxicity associated with Cas9 protein expression. Results showed differential levels of Cas9 toxicity among the bacteria and lower transformation efficiency for *cas9/dcas9* bearing plasmids compared to controls in general. This indicated lethal effect of Cas9/dCas9 expression. While *E. coli* and *S. typhimurium* seemed to tolerate Cas9/dCas9 fairly well, in GC rich microbes, *M. smegmatis, X. campestris* and *D. radiodurans*, Cas9/dCas9 associated toxicity was acute.

## Introduction

Recombinant DNA technology together with High Throughput Sequencing in recent times, has allowed us to harvest a large amount of genetic information from the microbial world. The technologies have been used extensively to find out which genes determine how microbes, grow, travel, starve, cause diseases, ward of predators and even die. This information is especially important for studying pathogenic bacteria, bacteria of industrial importance and ones with special stress tolerance abilities. Perturbing the normal functioning of the genome has emerged as the best method to probe function and dynamics of individual genes.

Discovery of the CRISPR -Cas viral defence systems opened up another novel and efficient tool box for genome editing, gene silencing, targeted gene methylation, etc. in all kinds of organisms from bacteria to humans [1][2][3][4]. The ease and efficiency of the system has made it an extremely popular *go-to* system for various applications. The system has further had widespread applications in metabolic engineering of bacteria as it allows easy programming and multiplexing [5][6]. Among the CRISPR-Cas systems, the Cas9 system from *Streptomyces pyogens* has gained popularity on account of being one of the earliest systems to be discovered and its simplicity of usage [7][8][9]. The system comprises a single protein, Cas9 and the sgRNA, which together can be easily employed to bring about a host of desired changes inside the cell of virtually any living being [10]. While the nucleoprotein, Cas9 itself has been extensively used for genome editing [7][8][11][9], its nuclease deficient variant, dCas9 has been useful for regulation of gene expression [3][1]. Systems employing the Cas9 variants have shown great promise for use in eukaryotic, particularly mammalian systems [11]. Attempts to use them in microbes have met with mixed success.

The Cas9/dCas9 and also Cas9 nickase systems were used successfully to probe gene function and cell dynamics in several microbes [7][2]. With the increasing use of Cas9, toxicity associated with Cas9 was noticed in certain microbes [12][13][14]. Further, toxicity was reported not only for the Cas9 protein, where non-specific nuclease activity would be expected to cause cell killing but also with the dCas9 protein, further compounding the problem. In a few cases, this problem could be circumvented by placing the *cas9/dcas9* genes under tight inducible control [12][14]. In a few microbes, even complete removal of promoter could not solve the toxicity [14]. The results indicated that different microbes have different threshold for tolerance towards *cas9/dcas9* expression which needs to be addressed for making the CRISPR-Cas9 system useful in such organisms.

In this study, we used appropriate plasmid systems to report *cas9/dCas9* mediated toxicity in five microbes, viz. *Escherichia coli, Salmonella typhimurium, Mycobacterium smegmatis, Xanthomonas campestris and Deinococcus radiodurans*. We investigated possible effect of methylation status of the host genome upon toxicity of *cas9/dcas9*. The study has also enabled comparisons of levels of toxicity imparted by *cas9* or *dCas9* in each organism and thus provides a comprehensive differential toxicity analysis in different groups of microbes.

## Materials and methods

### Bacterial strains, plasmids and growth conditions

*E. coli* and *S. typhimurium* cultures were grown in Luria Bertani medium (Tryptone Yeast extract and sodium chloride) at 37°C. *M. smegmatis* was grown in Middlebrook 7H9 with Tween 80 at 0.1% for with glycerol at 37°C. *D. radiodurans* was grown in Tryptone Glucose Yeast extract (TGY) broth at 32°C. *X. campestris* was grown in Luria Bertani medium at 28°C. All liquid cultures were grown with aeration at 180rpm, orbital shaking, The media were supplemented with Kanamycin, (50µg/ml for *E. coli* and 10 µg/ml for *M. smegmatis*) Carbenicillin (100 µg/ml for *E. coli*), Chloramphenicol (33 µg/ml for *E. coli* and 3 µg/ml for *D. radiodurans*), or Gentamycin (25 µg/ml), when necessary. Wherever required Anhydrotetracyclin (Atc) was added at a concentration of 1µM. The bacterial strains used are described in Table 1. The plasmid vectors used in this study are listed in Table 2.

**Table 1.** Bacterial strains used in the study.

| Strains used | Genotype | Source |
| --- | --- | --- |
| *E. coli* DH5alpha | F<sup>-</sup>, endA1, glnV44, thi-1, recA1, relA1, gyrA96, deoR, nupG, purB20, ϕ80dlacZΔM15, $\Delta$(lacZYA-argF)U169, hsdR17(rK-mK+), λ- | Lab collection |
| *E. coli* JM110 | rpsL, thr, leu, thi, lacY, galK, galT, ara, tonA, tsx, dam, dcm, glnV44, $\Delta$(lac-proAB), e14-, [F' traD36 proAB+ lacIq lacZΔM15], hsdR17, (rK-mK+) | Lab collection |
| *E. coli* JM109 | endA1, glnV44, thi-1, relA1, gyrA96, recA1, mcrB+, $\Delta$(lac-proAB), e14-, [F' traD36 proAB+ lacIq lacZΔM15], hsdR17, (rK-mK+) | Lab collection |
| *E. coli* GM2163 | F<sup>-</sup>, araC14, leuB6(Am), fhuA13, lacY1, tsx-78, glnX44(AS), galK2(Oc), galT22, λ-, mcrA0, dcm-6, hisG4(Oc), rfbC1, rpsL136(strR), dam-13::Tn9, xylA5, mtl-1, thiE1, mcrB9999, hsdR2 | Lab collection |
| *E. coli* Novablue | $\Delta$(srl-recA)30::Tn10(DE3), Tet<sup>r</sup> | Novagen |
| *Salmonella typhimurium* DA6192 | Wild type | Gift from Dr. Magnus Lundgren |
| *Mycobacterium smegmatis* MC<sup>2</sup> 155 | Wild type | Snapper et al. [32] |
| *Xanthomonas campestris* pv. Campestris str 8004 | Wild type | Lab collection |
| *Deinococcus radiodurans* R1 | Wild type (ATCC BAA 816) | Lennon et al. [33] |

**Table 2:**
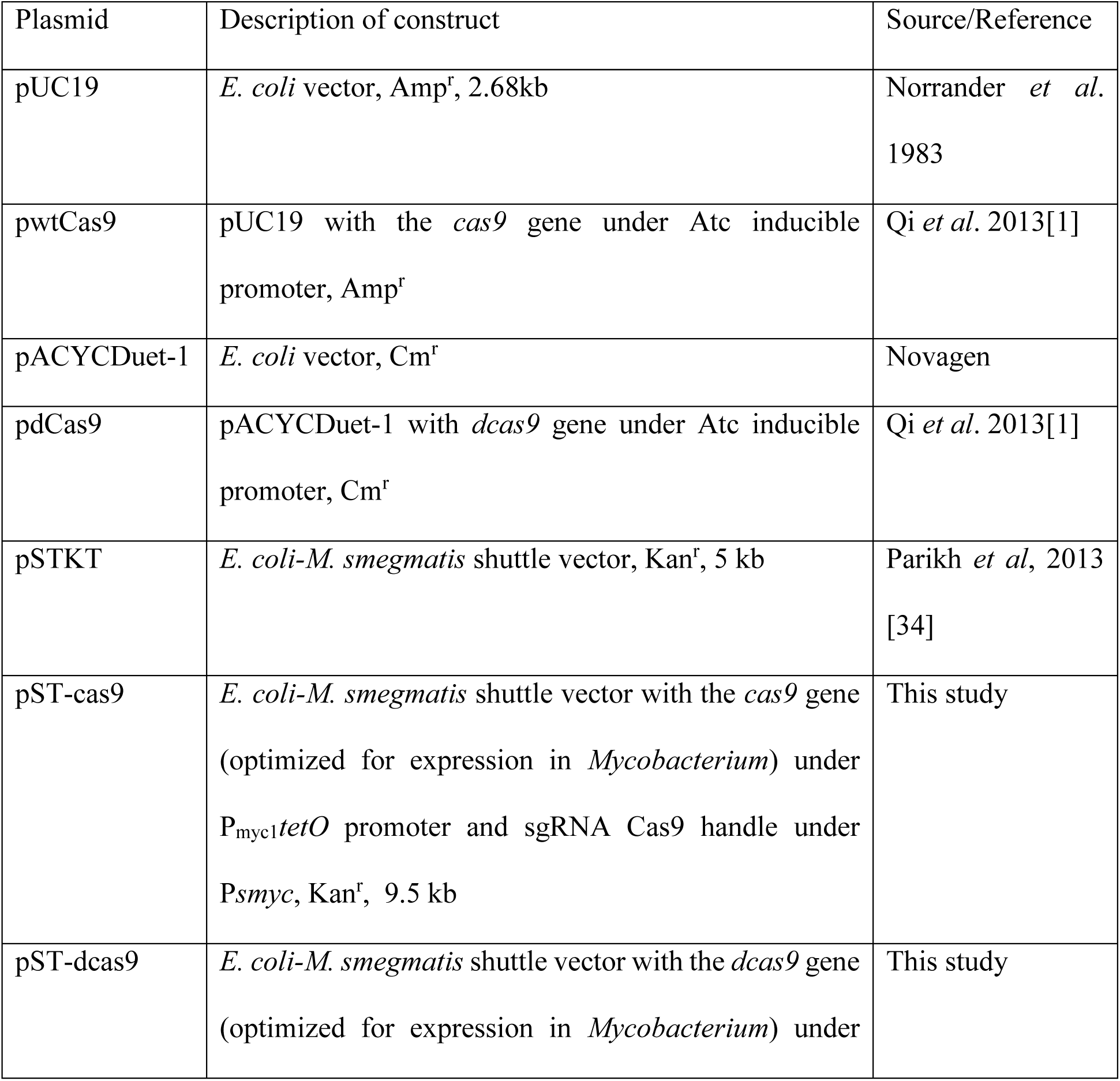
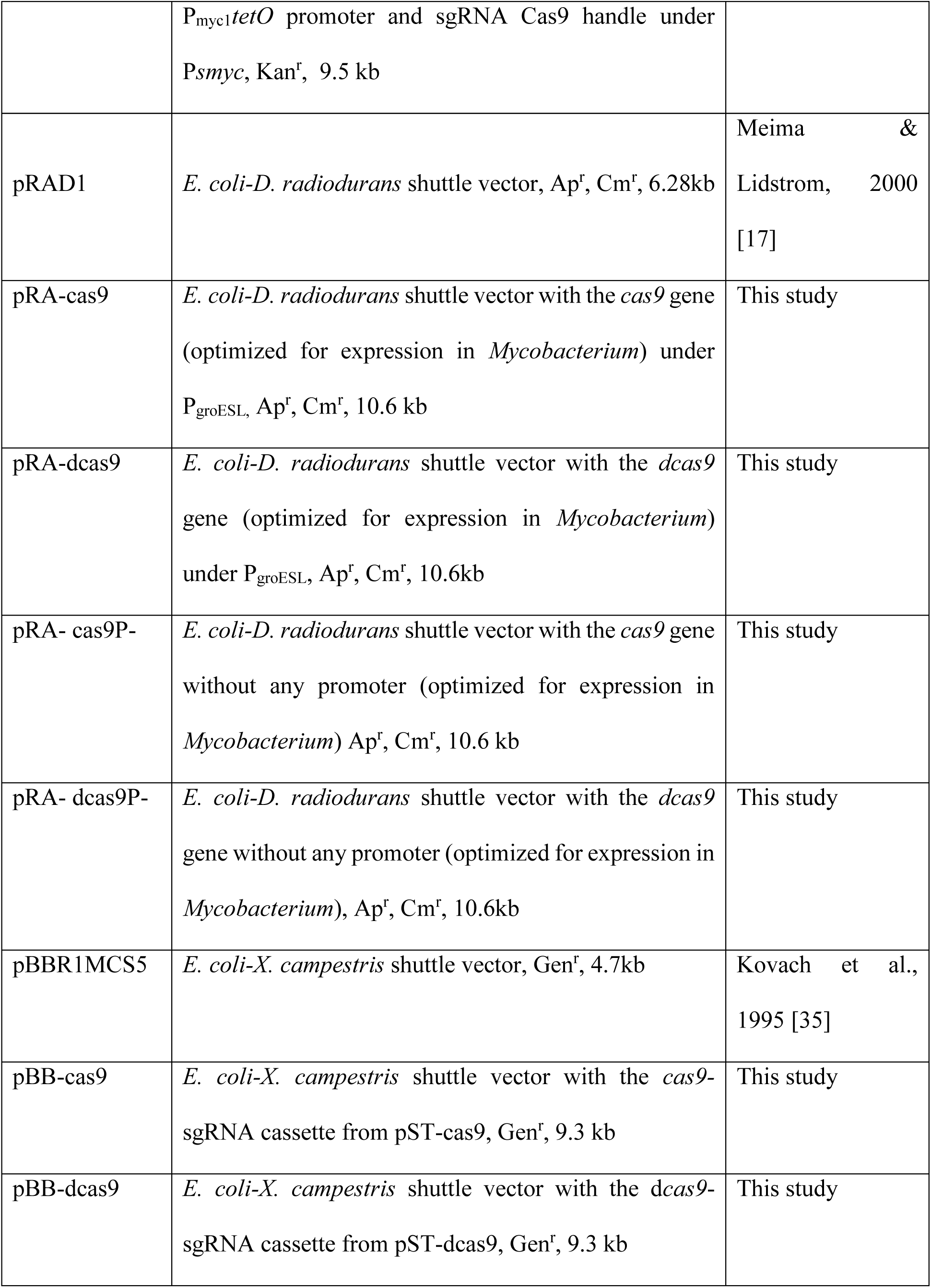
Plasmids used in this study

### Construction of the *cas9 – dCas9* expressing plasmids for use in *E. coli, S. typhimurium, M. smegmatis, X. campestris* and *D. radiodurans*

This study uses a host of plasmids that were either procured or constructed in various shuttle vectors under different promoters to suit their application in different microbes (Table 2). Of these, for *S. typhimurium* and *E. coli*, the wild type *cas9* from *Streptomyces pyogens* cloned in the pUC19 vector under the inducible promoter, PLtetO along with *dcas9* cloned in pACYduet under a similar inducible promoter were employed. These were procured as shown in Table 2.

#### Mycobacterium smegmatis

The gene for Cas9 was codon optimized for use in *Mycobacterium*. This was synthesized as a single fragment under P_myc1_*tetO*control while the Cas9 handle was placed under P_smyc_. The entire fragment was cloned into the KpnI -HindIII site of the multicopy vector, pSTKT to generate pST-cas9. Further the nuclease deficient mutant for Cas9 was generated by Gibson cloning [15] by generating the mutations D10A and H840A[16] The list of oligo primers used is given in Supplementary table1. The dCas9 thus generated, was similarly cloned into pSTKT to generate, pST-dcas9.

Results from the Graphical Codon Analyzer showed that the codon frequency in the *M. smegmatis-*optimized *cas9/dcas9* sequence matched well with codon usages in *D. radiodurans* and *X. campestris* (Supplementary Figures 1 & 2).

#### Xanthomonas campestris

The *cas9/das9* genes, optimized for *M. smegmatis* expression along with the promoters and sgRNA was cut out as a KpnI-HindIII cassette from pST-cas9 or pST-dcas9 and cloned into pBBR1MCS5 to generate, pBB-Cas9 and pBB-dCas9 respectively. Though the *cas9/dcas9* was under the inducible promoter, P_myc_*tetO* in *X. campestris*, due to absence of a Tet repressor, one would expect constitutive expression of the genes in this organism.

#### Deinococcus radiodurans

The open reading frames coding for Cas9/dCas9 optimized for *M. smegmatis* was cut out from pST-cas9 or pST-dcas9 using NdeI-BamHI restriction digestion and cloned into pRAD1 under control of the P_groESL_ to generate, pRA-Cas9 and pRA-dCas9 respectively. To generate suitable controls for transformation efficiency, the plasmids were also cloned without any promoter, pRA-Cas9P- and pRA-dCas9P-.

### Transformation of plasmid

*E. coli* and *D. radiodurans* were transformed into cells made competent using CaCl_2_ as described before [17]. *S. typhimurium, M. smegmatis* and *X. campestris* were transformed by electroporation. The details for electroporation have been given in Table 3.

**Table 3.** Electroporation conditions

| Organism | Electroporation apparatus | Voltage applied | Recovery conditions |
| --- | --- | --- | --- |
| *S. typhimurium* | Eppendorf eporator | 2.5kV | Luria Bertani broth at 37°C for 1h under shaking |
| *M. smegmatis* | Biorad Total Systems | 2.5kV | Middlebrook 7H9 with glycerol and 0.1% Tween 80 at 37°C for 2h under shaking |
| *X. campestris* | Eppendorf eporator | 1.4kV | Luria Bertani broth at 28°C for 4h under shaking |

### Expression of Cas9 and dCas9 in *E. coli*

Expression of Cas9 and dCas9 proteins was determined by separation of protein extracts of *E. coli* strains bearing the pRAD1, or pRA-cas9 and pRA-dcas9 by electrophoresis on a 10% denaturing polyacrylamide gel. The proteins were stained using Commassie Brilliant blue for visualization. The proteins were transferred to a PVDF membrane followed by incubation with Anti-Cas9 antibody conjugated to FITC (Sigma Aldrich). Anti-mouse secondary antibody conjugated to Alkaline phosphatase was used to develop the blot with NBT-BCIP (nitro-blue tetrazolium and 5-bromo-4-chloro-3’-indolyphosphate).

## Results

### Transformation efficiencies (TE) of plasmids bearing *cas9* in enterobacteria

In *S. typhimurium*, number of transformants recovered with pwtCas9 on Atc selection plates was half the number recovered in the absence of Atc. In Novablue strain, no transformants could be recovered on induction with Atc (Fig. 1a). In *E. coli* DH5 alpha, TE remained unaffected on Atc induction. However, Atc induction did result in smaller colony size of transformants in *E. coli* DH5alpha as well as *S. typhimurium* compared to uninduced culture carrying pwtCas9 but not pUC19 (Fig. 1c). This indicated that upon induction, toxicity also manifested as reduction in growth. Cas9 associated toxicity was therefore observed in *S. typhimurium* as well as *E. coli* Novablue, with a very pronounced effect in the latter.

**Fig. 1.**
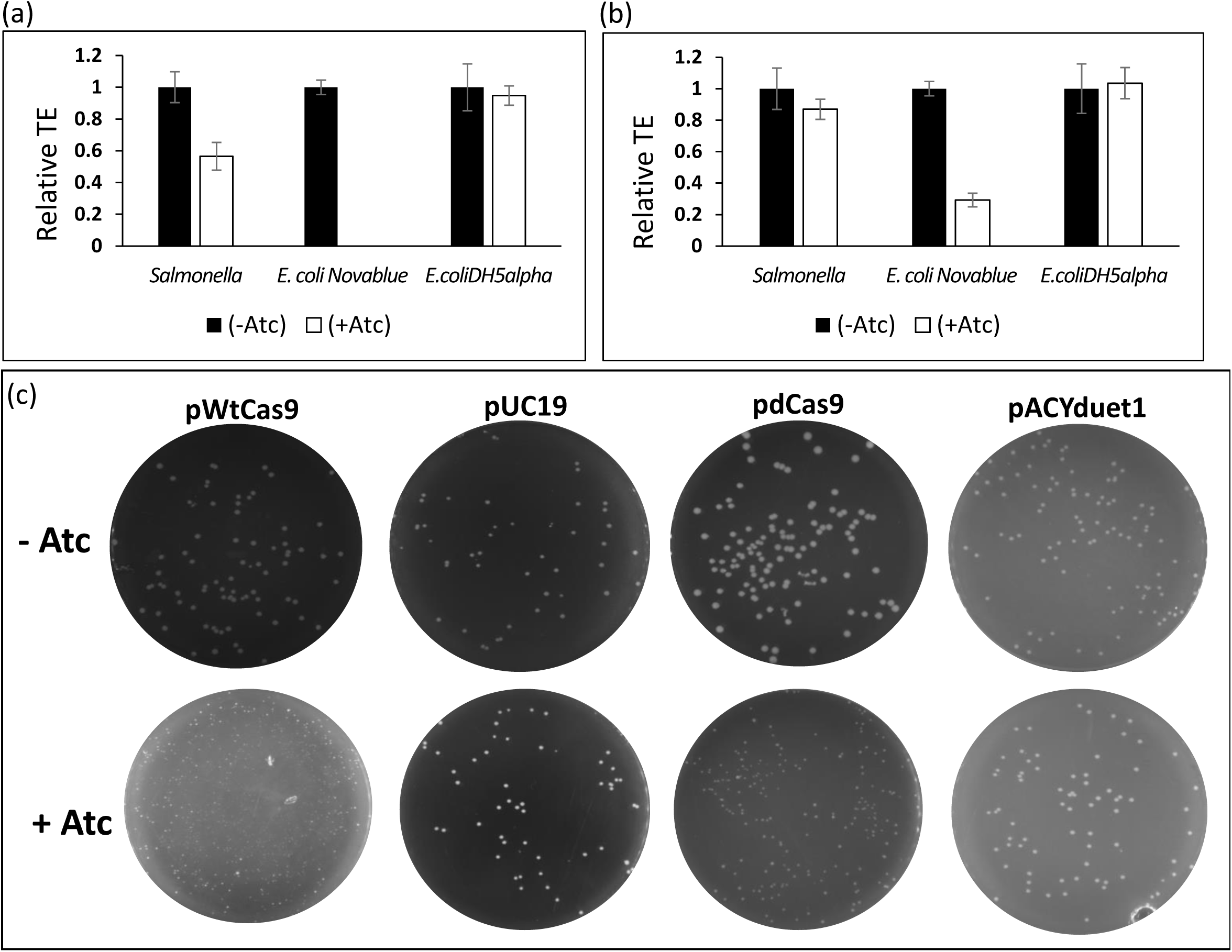
Transformation efficiency of plasmids bearing *cas9* (a) or *dcas9* (b) relative to the absence of induction with anhydrotetracycline in S. *typhimurium, E. coli* Novablue and *E. coli* DH5alpha. Transformation was done by electroporation for *S. typhimurium* and by CaCl_2_ method for *E. coli* cells. Transformants were selected on antibiotic selection plates and colony forming units were enumerated to determine transformation efficiency. (c) Reduction of colony size on induction with Atc as observed for *S. typhimurium.* Results are from experiments that were repeated three times each.

### Transformation efficiencies of plasmids bearing *dcas9* in enterobacteria

In *S. typhimurium*, number of colonies recovered with pdCas9 on Atc induction was comparable to that in the absence of induction. However, as in case of pwtCas9, the size of the colonies was smaller on Atc induction of dCas9 expression (Fig. 1c). In Novablue, TE with pdCas9 decreased around three fold on Atc induction, while it was unaffected when pACYCDuet-1 was used (Fig. 1b). In *E. coli* DH5 alpha, TE was unaffected on Atc induction. The results indicate that dCas9 mediated toxicity was highest in *E. coli* Novablue followed by*S. typhimurium*, while *E. coli* DH5 alpha tolerated dCas9 expression well.

### Transformation efficiencies of plasmids bearing *cas9/dcas9* in *M. smegmatis* and *X. campestris*

In *X. campestris*, no transformants for pBB-cas9 or pBB-dcas9 could be recovered, while on average, a TE of 1.5×10^3^ CFU/ µg of DNA could be obtained with the vector control (Fig. 2a). Rarely a few transformants were recovered only in the case of dCas9 which did not grow subsequently in liquid medium.

**Fig. 2.**
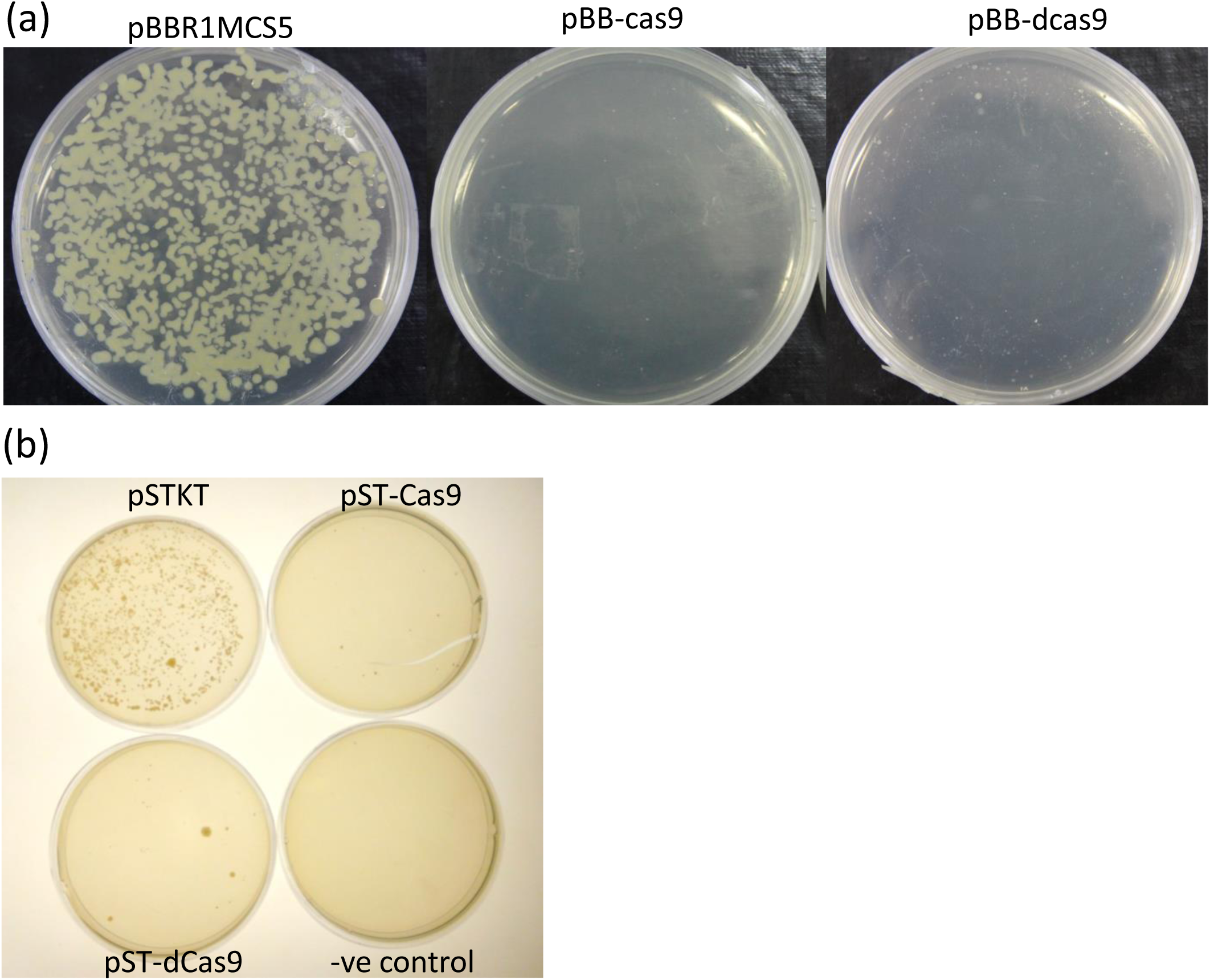
Transformation of cas9/dcas9 bearing plasmids in *X. campestris* (a) and *M. smegmatis* (b). Plasmids bearing *cas9/dcas9* were electroporated along with empty vector controls into *X. campestris* and *M. smegmatis*. The cells were spread on antibiotic selection plates and allowed to grow.

In *M. smegmatis* also, transformation of pST-cas9 and pST-dcas9 plasmids yielded no transformants even when no Atc was added to the selection plates, while in empty vector control, a TE of 7.6×10^3^ CFU/µg DNA was obtained (Fig. 2b). Here, too, rarely small colonies of transformants could be recovered on selection plates that failed to grow in liquid culture. Both these organisms, therefore displayed acute toxicity to both Cas9 as well as dCas9.

### Transformation efficiencies of plasmids bearing *cas9/dcas9* in *D. radiodurans*

On transforming *D. radiodurans* with pRA-Cas9 and pRA-dCas9 plasmids, no colonies could be recovered, while with control plasmids, an average TE of 1.5 × 10^3^CFU/μg DNA could be obtained (Fig. 3a). Even when *cas9/dcas9* plasmids without the P_groESL_ promoter were used, transformants could not be recovered. To increase the overall transformation efficiency in this organism, the plasmids were passaged through a dam^−^/dcm^−^ strain, *E. coli* JM110. Plasmids isolated from this *E. coli* strain when transformed into *D. radiodurans* resulted in a higher transformation efficiency of control plasmid, pRAD1 (average of 2.3 ×10^4^ CFU/μg DNA) (Fig. 3b & c). Importantly, at this transformation efficiency, *cas9/dcas9* bearing transformants could be recovered at an efficiency of 1.7×10^2^ and 2.9×10^2^ CFU/µg DNA respectively which was 100 fold lower than that for pRAD1 (Fig. 3c). Transformants were also recovered at an efficiency of 2×10^3^ and 2.5×10^3^ CFU/ μg DNA with *cas9* and *dcas9* bearing plasmids in the absence of a promoter (Fig. 3b & c). The results indicate that toxicity of Cas9/dCas9 in *D. radiodurans* is moderate.

**Fig. 3.**
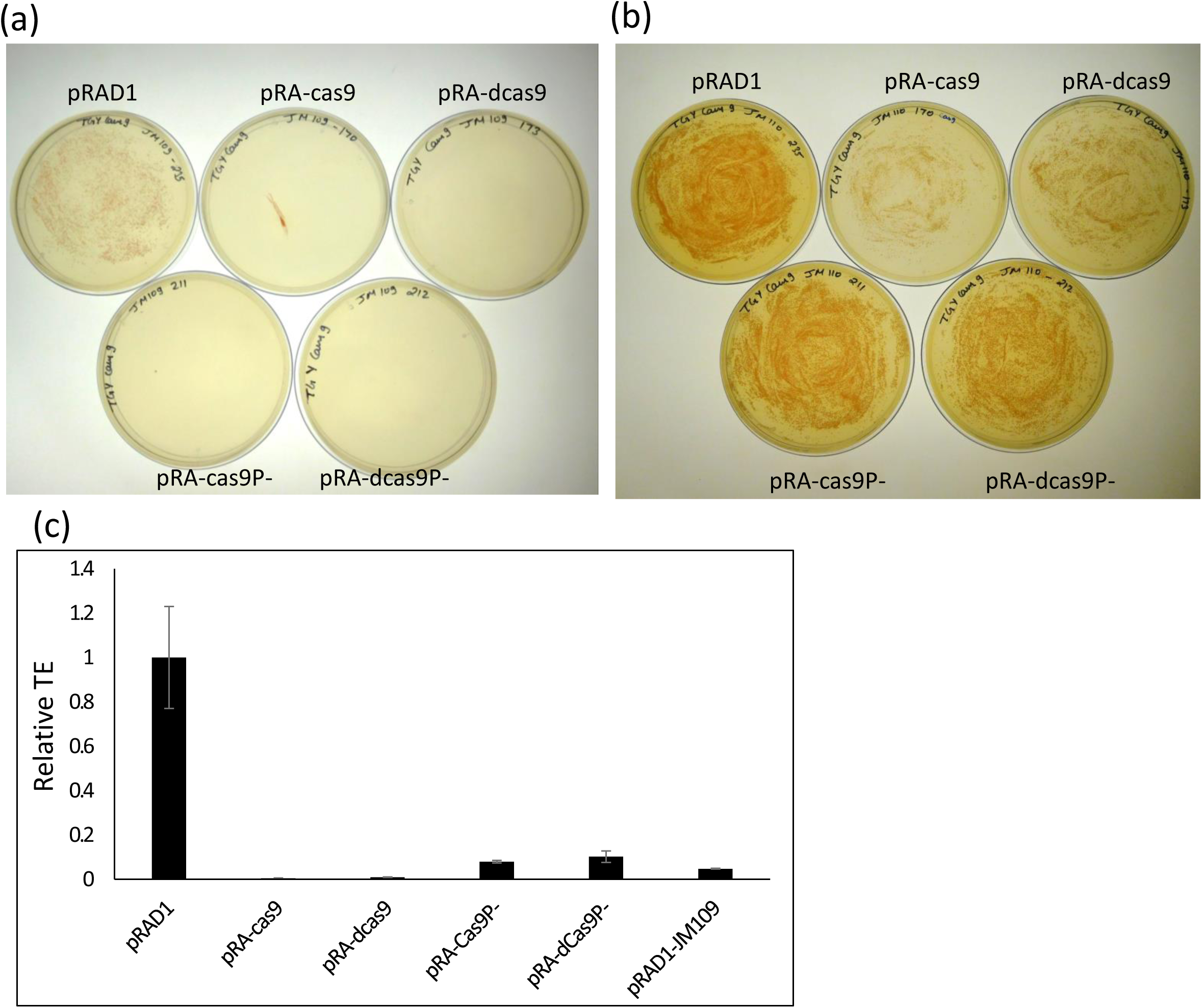
Transformation of plasmids pRAD1, and pRAD1 bearing cas9 and dcas9 with (pRA-cas9 and pRA-dcas9) or without promoter (pRA-cas9P- and pRA-dcas9P-) in *D. radiodurans*. Plasmids isolated from JM109(a) or JM110(b) strains of *E. coli* were used to transform *D. radiodurans.* Cells were plated on chloramphenicol selection plates. (c) Transformation efficiency of plasmids as determined from CFUs enumerated on antibiotic selection plates where colonies could be recovered. Results are from experiments repeated three times.

### Effect of DNA methylation on Cas9/dCas9 toxicities

As Cas9 is a nuclease and dCas9 retains the DNA binding property, the effect of methylation of genomic DNA on Cas9/dCas9 mediated toxicity was analysed. Two strains of *E. coli* that were dam^+^/dcm^+^, DH5alpha and JM109 and two strains that were dam^−^/dcm^−^, JM110 and GM2163 were employed. Plasmids, pRA-cas9 and pRA-dcas9 where Cas9/dCas9 is expressed from a strong constitutive promoter P_groESL_ were transformed into each of these *E. coli* strains. To incorporate size control for such plasmids, pRA-cas9P- or pRA-dcas9P-were also transformed into all *E. coli* strains. Results did not show a marked effect for DNA methylation in determining Cas9/dCas9 based toxicity (Fig. 4a). However, the fraction of transformants obtained with *cas9/dcas9* bearing plasmids compared to promoterless plasmids was marginally fewer in dam^−^/dcm^−^ strains compared to dam^+^/dcm^+^ strains (Fig. 4a). To determine expression of Cas9/dCas9 in each strain of *E. coli*, cell extracts were separated on a SDS-PAGE gel by electrophoresis and visualized by Coomassie staining and Western blot. Cas9/dCas9 expression was seen in all the *E. coli* strains carrying *cas9/dcas9* bearing plasmids as a 160 kDa band that was absent in empty vector control (Fig. 4b &c). The levels of Cas9/dCas9 expression was higher in dam^+^/dcm^+^ strains compared to that in dam^−^/dcm^−^ strains, with the highest expression in DH5alpha and lowest in GM2163 strain (Fig. 4b & c). The results indicate marginally higher toxicity of Cas9/dCas9 dam^−^/dcm^−^ strains compared to dam^+^/dcm^+^ strains despite lower protein expression.

**Fig.4.**
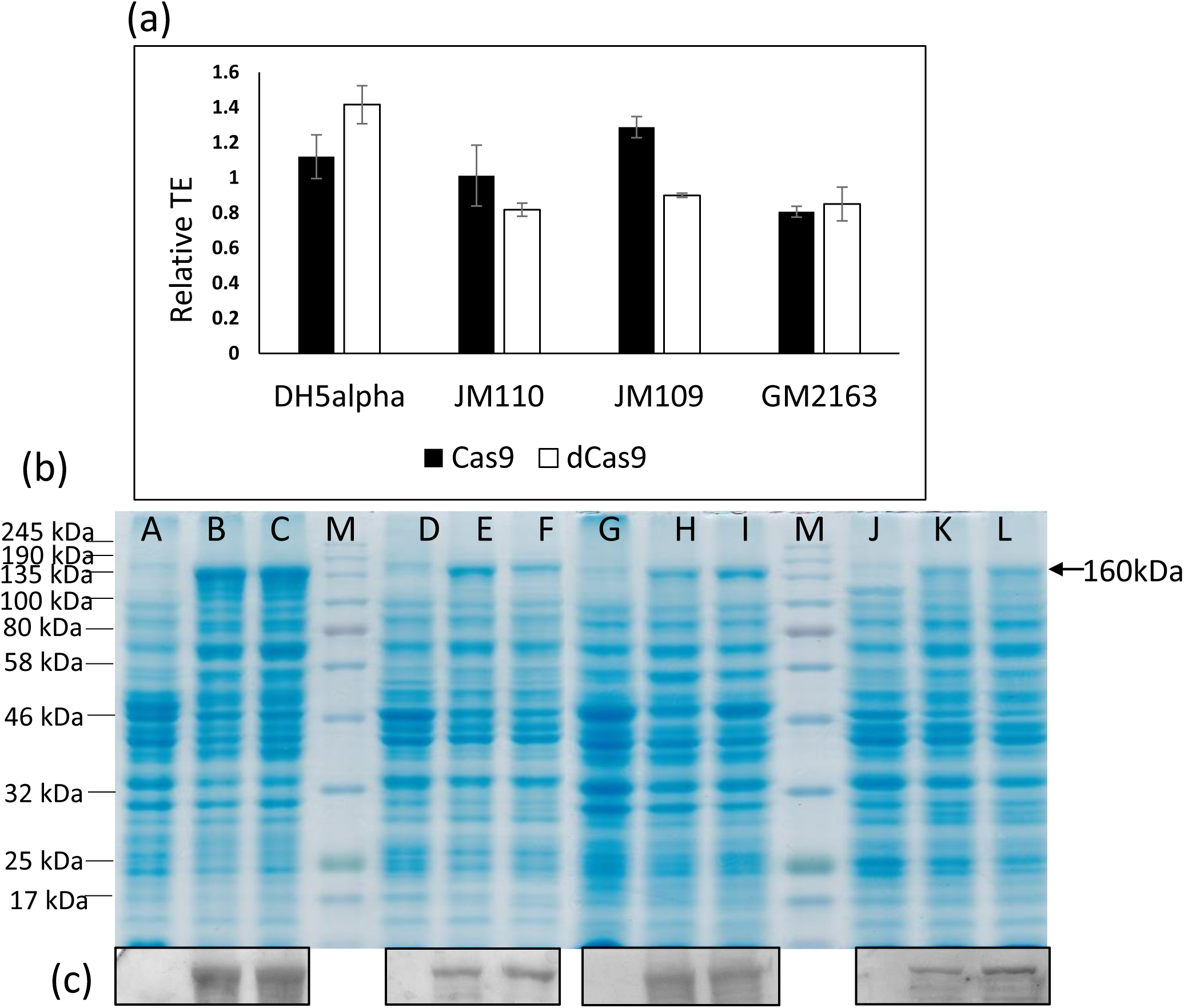
Cas9/dCas9 associated toxicity in *E. coli* strains that are dam^+^/dcm^+^ or dam^−^/dcm^−^.Four *E. coli* strains were transformed with pRAD1 or pRA-Cas9 or pRA-dCas9 and transformants were elected on carbenicillin selection plates. The CFU were enumerated to determine TE (a). Results are from experiments repeated four times. (b) Protein extracts from *E. coli* strains, DH5alpha (Lanes A, B,C), JM110 (Lanes D, E, F), JM109 (Lanes G, H, I) and GM2163 (Lanes J, K,L) carrying pRAD1 (Lanes A, D, G and J) or pRA-Cas9 (Lanes B, E, H and K) or pRA-dCas9 (Lanes C, F, I and L) were separated by gel electrophoresis and stained with Coomassie Brilliant Blue (b) or developed for Western blot using Anti-Cas9 antibody (c).

## Discussion

Cas9/dCas9 CRISPR system has enormous potential to be applied in a wide variety of organisms but the challenge in utilizing its full potential is the toxicity associated with it. Cas9/dCas9 associated toxicities have been found in many microbes such as *Synechococcus elongates* UTEX [14], *E. coli* [18][19], *M. smegmatis* [12], *Chlamydomonas reinhardtii* [20] *Corynebacterium glutamicum* [6] etc. The reason for toxicity of Cas9/dCas9 in microbes has been alternating between the obvious and the mysterious ever since development of this technology. Early reports put down toxicity of the Cas9 protein to possible non-specific nuclease activity. But similar toxicity with dCas9 required the theory to be revised. In *M. smegmatis* it was reported that dCas9 causes proteotoxicity that sensitizes the bacteria to stress [12]. The last two years have provided more insights on *cas9/dcas9* toxicity. Overall, it emerged that by keeping the expression levels low or only transiently expressing these proteins, toxicity could be avoided [14][12]. The determinants of Cas9 toxicity however remained elusive.

In this study, we bring forth toxicity data for Cas9/dCas9 in five different bacteria, two of which (*M. smegmatis* and *E. coli*) are standard model organisms, two are pathogens, (*X. campestris* and *S. typhimurium*) and one shows phenomenal stress tolerance to radiation (*D. radiodurans*). Comparison of transformation efficiencies obtained with *cas9/dcas9* bearing plasmids against empty vector or promoter-less controls, or on induction of protein expression have been interpreted to reflect Cas9/dCas9 mediated toxicity. In all cases, the choice of plasmids and promoters was guided by well-established expression systems for the respective organism. Therefore, though toxicities of Cas9/dCas9 are discussed across the five microbes in this study, they are not directly comparable in absence of a way to normalize expression levels. Wild type versions of *cas9/dcas9* were employed in experiments involving *E. coli* and *S. typhimurium* under an inducible promoter, while for *M. smegmatis, X. campestris, D. radiodurans*, and also *E. coli* for certain experiments, the genes were codon optimized for use in *M. smegmatis* that also fulfilled codon usages for *X. campestris* and *D. radiodurans.*

In all the organisms tested, generally, lower transformation efficiencies were obtained with *cas9/dcas9* bearing plasmids compared to empty vector control or the promoter-less versions of plasmids or in absence of inducer for Cas9/dCas9 expression. Acute Cas9 mediated toxicity was observed in *X. campestris, M. smegmatis* while *D. radiodurans* exhibited moderate toxicity. In *M. smegmatis*, toxicity was severe even in the absence of induction, while in others, observed toxicities were due to *cas9/dcas9* expression that was driven constitutively. All earlier studies where dCas9 systems were used in *M. smegmatis* involved use of integrative plasmids with tight control on expression [21][22][12] due to problems associated with dCas9 toxicity in *M. smegmatis*. A direct comparison between Cas9 and dCas9 toxicity was possible only in the systems where the same plasmid and promoter systems were utilized (*D. radiodurans, X. campestris, M. smegmatis* and *E. coli* where pRAD1 based systems were employed) and the results showed that the toxicity due to Cas9 and dCas9 was comparable.

Several studies showed that Cas9/dCas9 toxicity become evident at high levels of expression of the proteins even in *E. coli* [18][13]. Further a bad seed effect describing effect of certain sgRNAs known to not target essential genes was also described in *E. coli* [18]. One study reported changes in expression levels of several genes and cell morphology in *E. coli* at high levels of dCas9 expression [13]. In our study too, using either the wild type version or the codon optimized *cas9/dcas9* caused a certain degree of toxicity with both inducible and constitutive systems *in E. coli*. Surprisingly, the different strains employed, showed large variations in their response to Cas9/dCas9 expression. DH5alpha seemed to be the most robust strain that remained minimally affected with both versions of the *cas9/dcas9* genes, in spite of high expression of the protein. But, even this strain, Atc induction of Cas9 expression resulted in a reduction in colony size but not TE. This was not the case with *dcas9*, where the transformation efficiency as well as colony size was unaffected on induction of the gene. JM109 also seemed to tolerate Cas9/dCas9 expression relatively well when tested with constitutively expressed genes. The Novablue strain of *E. coli* showed moderate toxicity with the two genes, but on induction of Cas9, no transformants could be recovered. The results are useful while choosing between *E. coli* strains for applications involving Cas9 and its variants.

In the last two years, investigations have indicated that Cas9/dCas9 toxicity is perhaps due to non-specific binding to NGG sequences in the genome and the unwinding of the genome for PAM searching [19]. The threshold concentration of dCas9 at which toxicity just appears in *E. coli* could be increased by abolishing the PAM binding property in dCas9, lending credibility to this theory. Further, earlier reports have shown high-affinity non-specific DNA binding by the Cas9 in the absence of sgRNA [23].

This would also imply that a determinant of Cas9/dCas9 toxicity in an organism is amenability of its chromosome to binding by such proteins. This would in turn almost certainly depend on the GC content of the organism influencing PAM density on genome, the fraction of the genome that is transcriptionally active and perhaps its epigenetic status. The microbes that showed acute toxicity towards Cas9/dCas9 in this study are all GC rich, resulting in higher occurrence of NGG in the genome leading to binding of the chromosome at higher density resulting in disruption of normal DNA metabolism. This explains why an organism such as *D. radiodurans* which is known for its ability to repair DNA damage, would also suffer from Cas9/dCas9 related toxicity. Another observation from this study was that the toxicity in all strains was marginally higher for *cas9* bearing plasmids than *dCas9* bearing plasmids. This is again expected considering that Cas9 might also exert a non-specific, sgRNA independent cleavage of the chromosome at a low frequency, that would be absent in *dcas9* expressing strains.

In eucaryotes and other cell lines, the chromosome is more tightly packed and organized, that may lead to low accessibility of the DNA for non-specific Cas9 interactions. Further even if such interactions do occur at a low frequency, it may be buffered by the presence of a higher percentage of ‘junk’ DNA than in procaryotes, therefore not interfering with DNA metabolism sufficiently to cause toxicity. Nevertheless, reports on definitive Cas9/dCas9 mediated toxicities in eucaryotes, especially single-celled organismssuch as *Toxoplasma gondii* [24], yeast [25], *Trichomonas vaginilis* [26], have begun to appear in literature. Plasmids carrying Cas9 have also shown toxicity in some cell lines where ribonucleoprotein delivery has improved viability [27].

This study also attempted to evaluate effect of host genome methylation on the toxicity of Cas9/dCas9. Since, dam^+^/dcm^+^ strains would carry methylated chromosomes, it is tempting to assume that this modification would mask the DNA to discourage non-specific Cas9/dCas9 binding and lead to lower toxicity levels. In dam^−^/dcm^−^ strains, a naked DNA would probably make non-specific Cas9/dCas9 binding easier. It is another question, whether the frequency of methylated sites on the *E. coli* chromosome in dam+/dcm+ strains would be enough to effect non-specific binding resulting in toxicity at all or not. Our results indicate marginally lower Cas9/dCas9 mediated toxicity in dam^−^/dcm^−^ strains despite lower expression of proteins. Therefore, the results are not sufficient to conclude no effect due to chromosomal methylation as far as Cas9/dCas9 binding is concerned. Earlier, it was shown that cleavage by Cas9 was unaffected by cpG methylation [28]. However, there are other studies which showed that there was a negative co-relation between off-target binding and DNA methylation [29]. It remains to be proven more rigorously, whether indeed bacterial methylation affects non-specific binding if at all, hence influencing toxicity and possibly off-target effects or whether it also affects targeting.

## Conclusion

In view of results from this study, Cas9 expression in microbes may need careful modulation to ensure effective applications in silencing and genome editing. Especially in complex systems such as metabolic engineering, Cas9/dCas9 toxicity may need evaluation, since multiple gene modifications may compound the toxicity problem. It may be useful to employ strategies such as use of temperature sensitive plasmids, integrative plasmids or low copy number plasmids, transient expression or direct use of the sgRNA-Cas9 nucleprotein complex by electroporation etc. to minimise the toxic effects of the protein. In addition, alternative CRISPR systems that maybe associated with lower toxicity such as TypeI (Cascade) [30] or Type V (Cpf1) [31] may be explored where specifically essential genes need to be probed.

## Supporting information

Primers for generating dCas9 by Gibson cloning

Codon usage results for Cas9 in D. radiodurans and X. campestris

## Acknowledgements

We would like to acknowledge Addgene for providing the plasmids, pwtCas9 and pdCAs9, Dr. Magnus Lundgren, Uppsala University for *S. typhimurium* DA6192 strain and Dr. Amit Singh, Indian Institute of Science, Bengaluru for *M. smegmatis.* We would like to thank Dr. Shyam Sunder Rangu, Dr. Bhakti Basu and Dr. Yogendra S. Rajpurohit for their inputs on the manuscript. We also acknowledge Dr. Hari S. Misra for constant help and support.

