## Supplementary material for "Determination of Cas9/dCas9 associated toxicity in microbes": Primers for generating dCas9 by Gibson cloning

Supplementary table 1. Primers used for construction of dCas9 mutants by Gibson cloning

| Primer | Sequence (5’ to 3’) |
| --- | --- |
| dCas91-f | atgcggaggaatcacttccatATGGACAAGAAGTACAGCATCGGCCTGGCCATCGGCACC-3’ |
| dCas91-r | atggcgtccaCGTCGTAGTCGCTCAGCCGGTTGATGTCCA |
| dCas92-f | gactacgacgTGGACGCCATCGTGCCGC |
| dCas92-r | cggccgctctagagatatcggatccTCAGTCGCCGCCCAGCTG |
