## Supplementary material for "Determination of Cas9/dCas9 associated toxicity in microbes": Codon usage results for Cas9 in D. radiodurans and X. campestris

www.gcua.de
created: 18.10.2019

Cas9
sequence derived from M. smegmatis

Codontable:
https://www.kazusa.or.jp/codon/cgi-bin/showcodon.cgi?species=1299&aa=1&style=N

Ordinate (y-axis): frequency <20% <10%

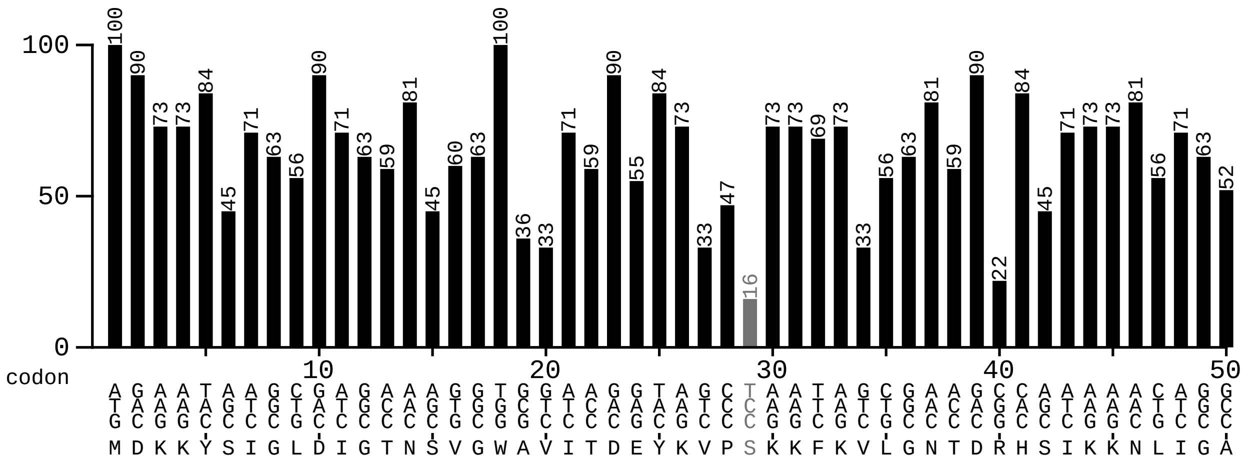


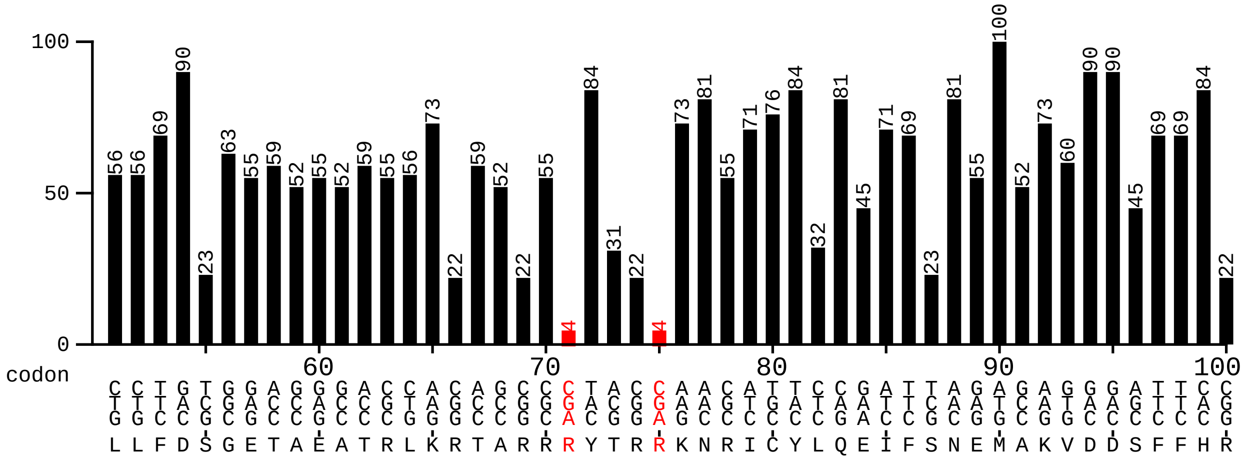


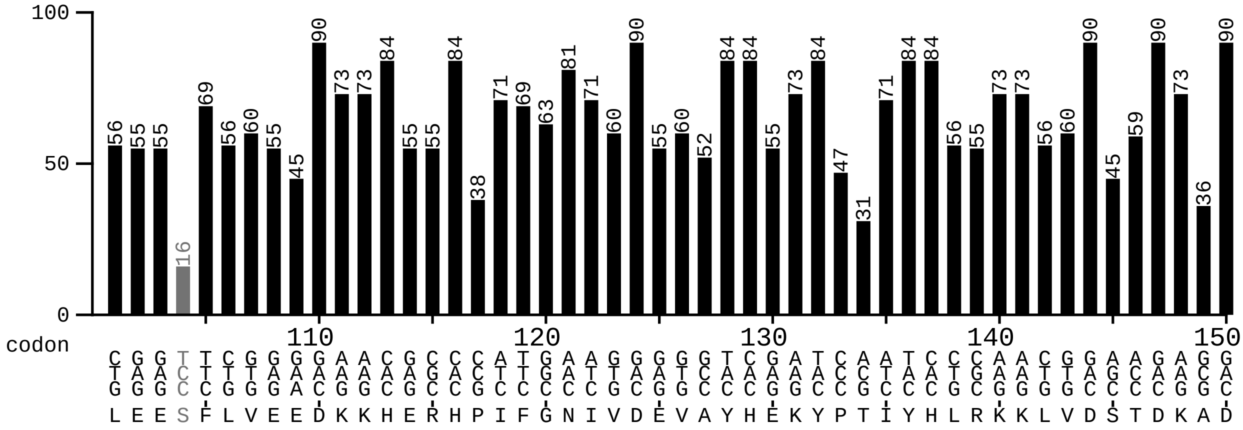

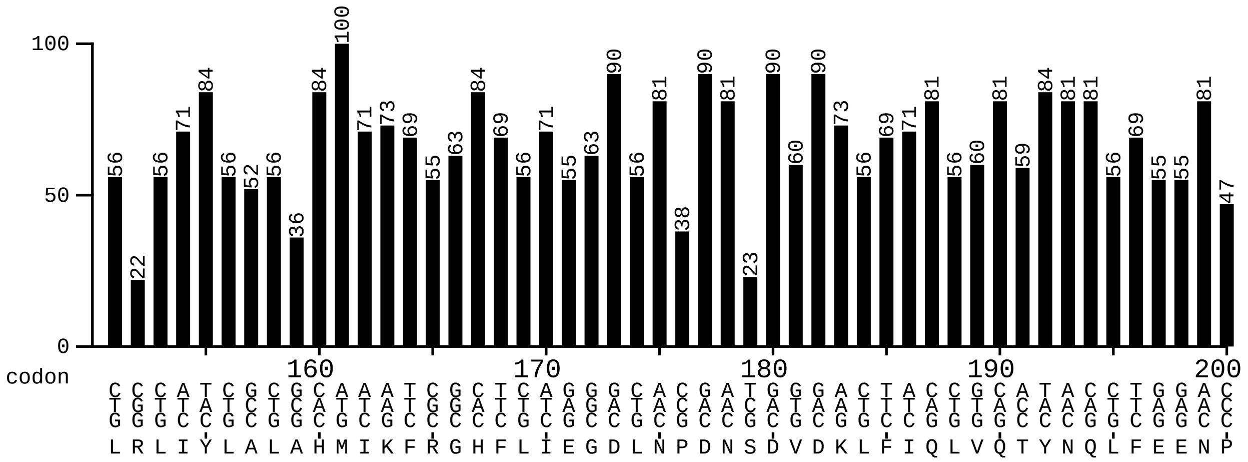
100
50
0
codon
160
170
180
190
200

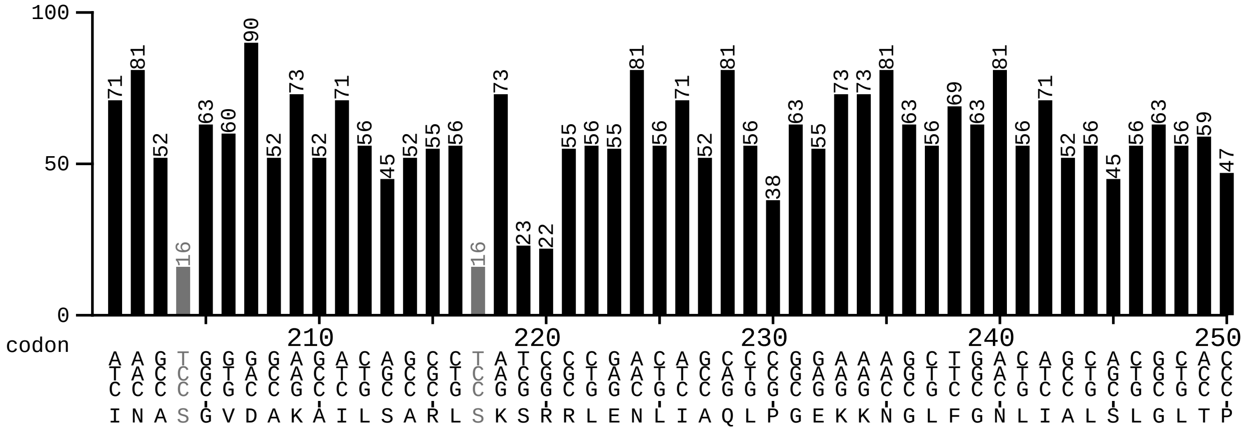
100
50
0
codon
210
220
230
240
250

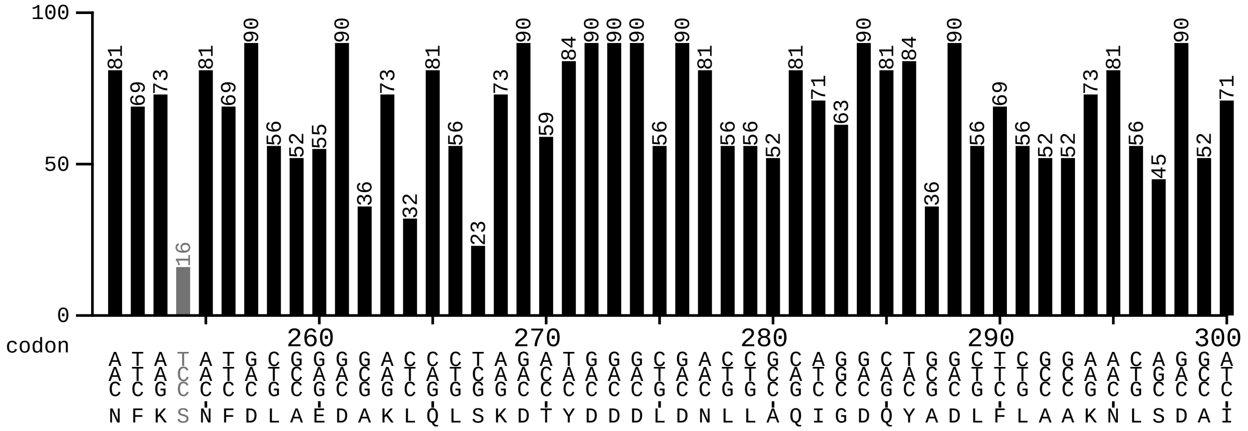
100
50
0
codon
260
270
280
290
300

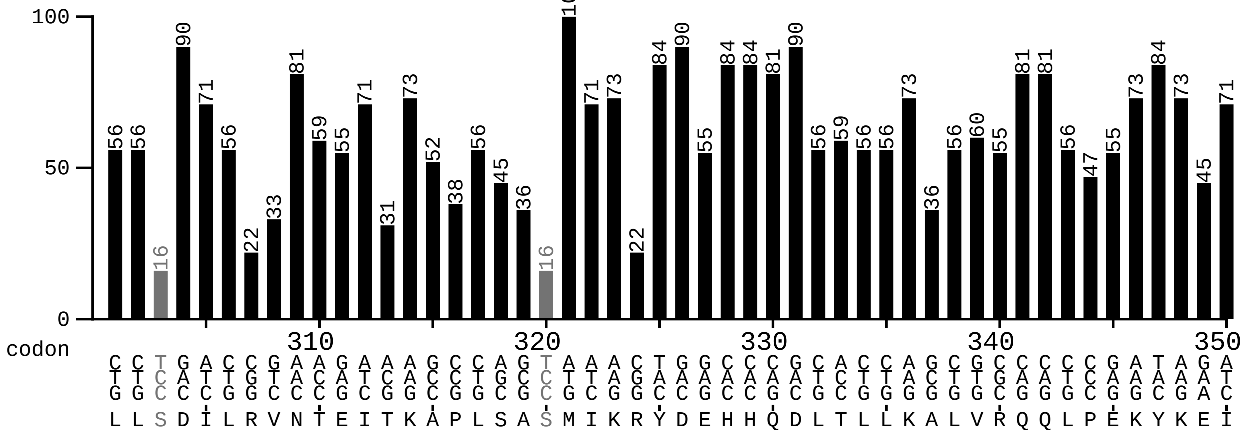
100
50
0
codon
310
320
330
340
350

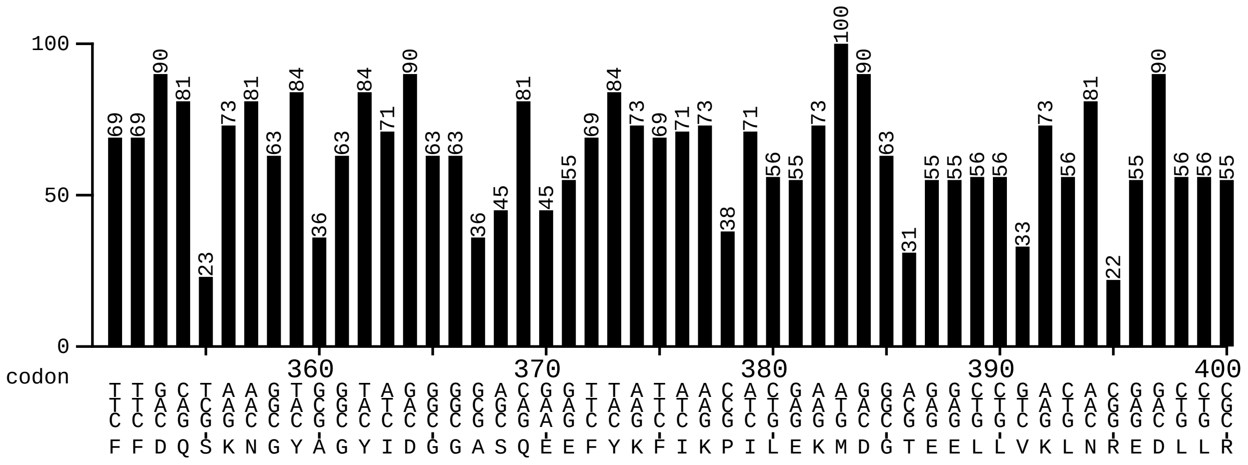
100
50
0
codon
360 370 380 390 400
F F D Q S K N G Y A G Y I D G G A S Q E E F Y K F I K P I L E K M D G T E E L L V K L N R E D L L R

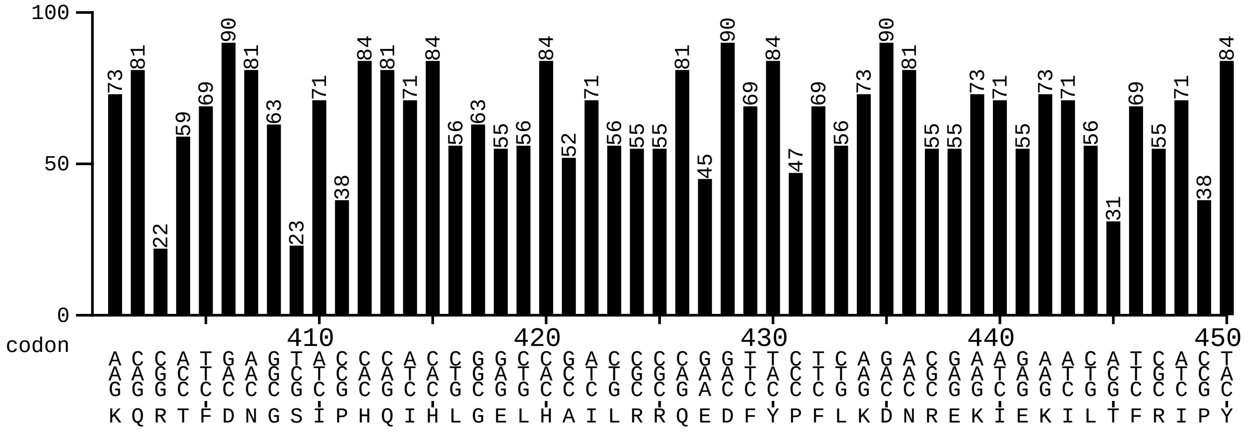
100
50
0
codon
410 420 430 440 450
K Q R T F D N G S I P H Q I H L G E L H A I L R R Q E D F Y P F L K D N R E K I E K I L T F R I P Y

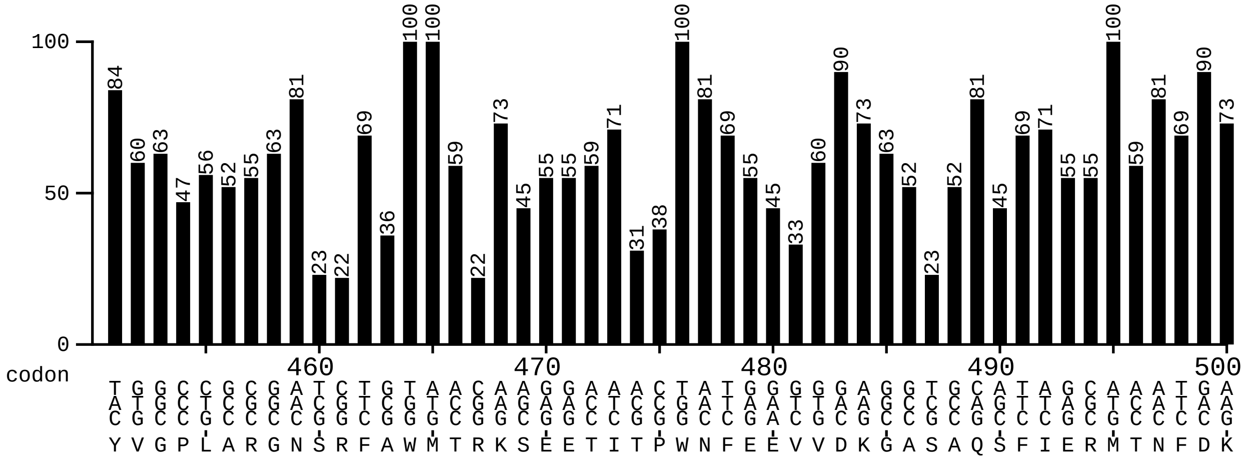
100
50
0
codon
460 470 480 490 500
Y V G P L A R G N S R F A W M T R K S E E T I T P W N F E E V V D K G A S A Q S F I E R M T N F D K

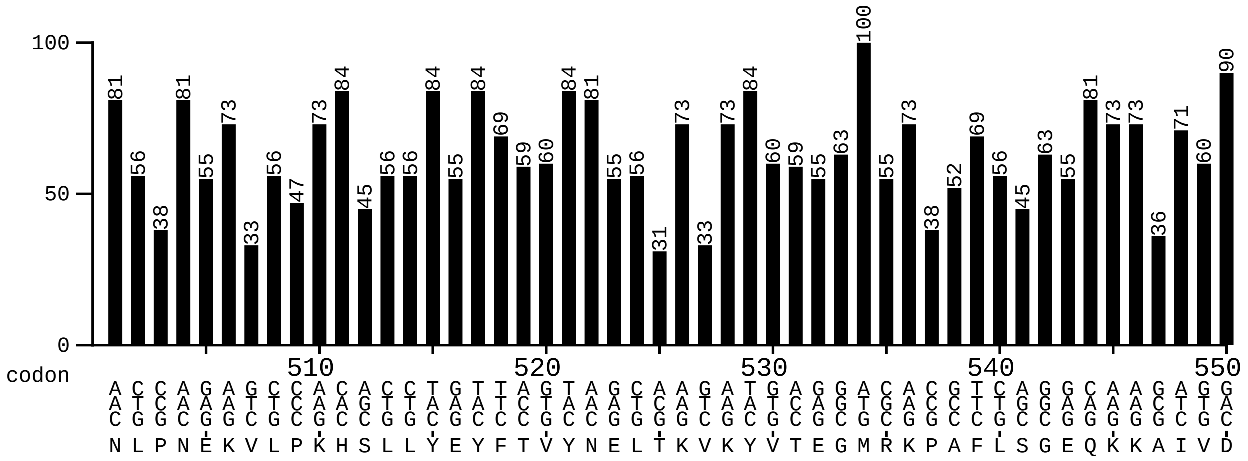
100
50
0
codon
510 520 530 540 550
N L P N E K V L P K H S L L Y E Y F T V Y N E L T K V K Y V T E G M R K P A F L S G E Q K K A I V D

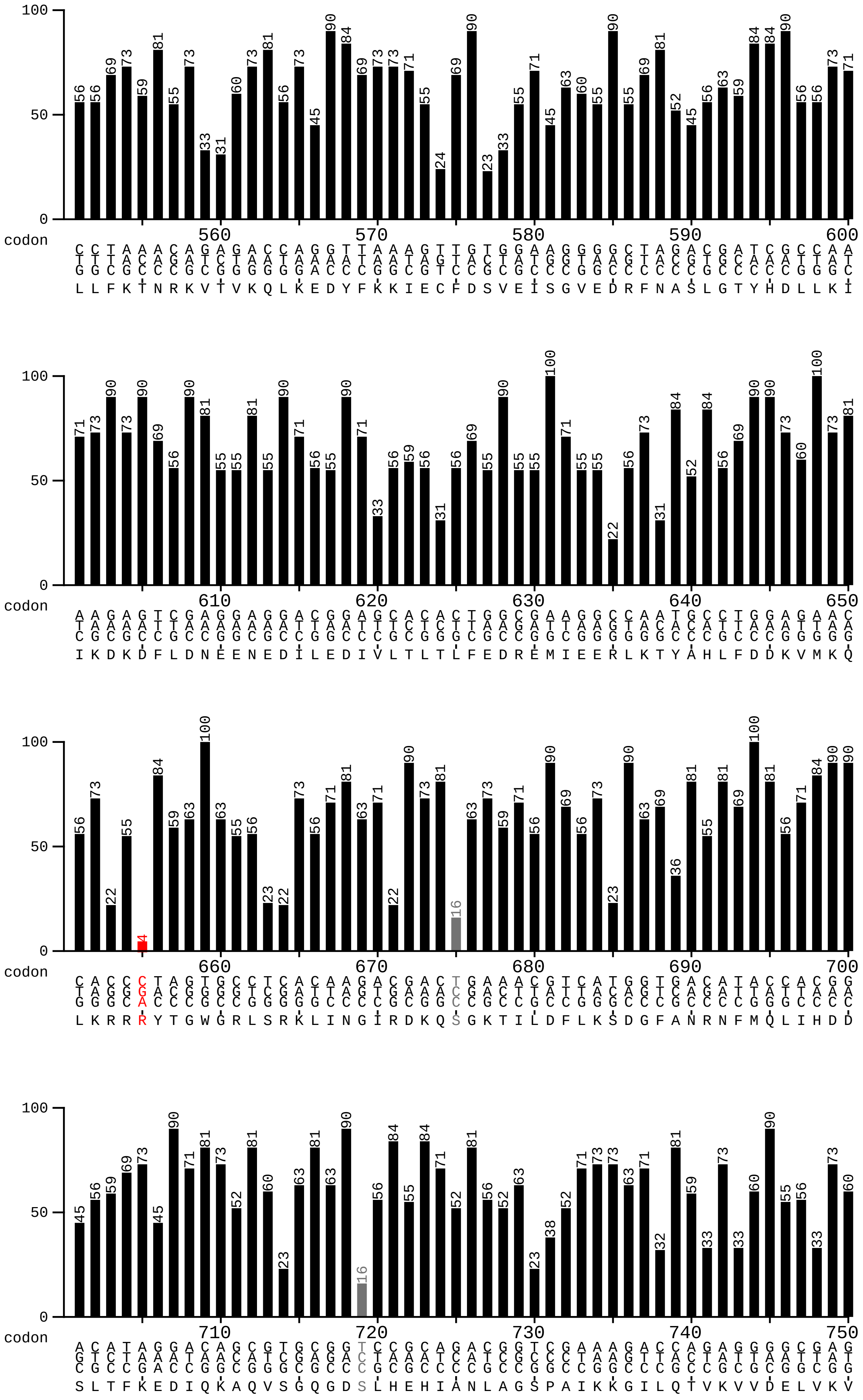
100
50
0
codon
560 570 580 590 600
L L F K T N R K V T V K Q L K E D Y F K K I E C F D S V E I S G V E D R F N A S L G T Y H D L L K I
610 620 630 640 650
I K D K D F L D N E E N E D I L E D I V L T L T L F E D R E M I E E R L K T Y A H L F D D K V M K Q
660 670 680 690 700
L K R R R Y T G W G R L S R K L I N G I R D K Q S G K T I L D F L K S D G F A N R N F M Q L I H D D
710 720 730 740 750
S L T F K E D I Q K A Q V S G Q G D S L H E H I A N L A G S P A I K K G I L Q T V K V V D E L V K V

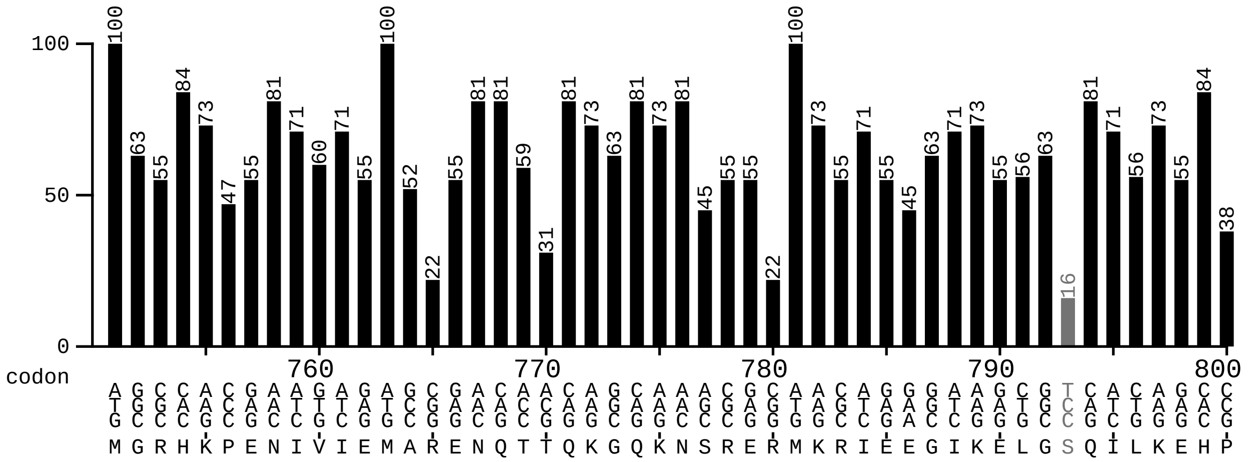
100
50
0
codon
760 770 780 790 800
M G R H K P E N I V I E M A R E N Q T T Q K G Q K N S R E R M K R I E E G I K E L G S Q I L K E H P

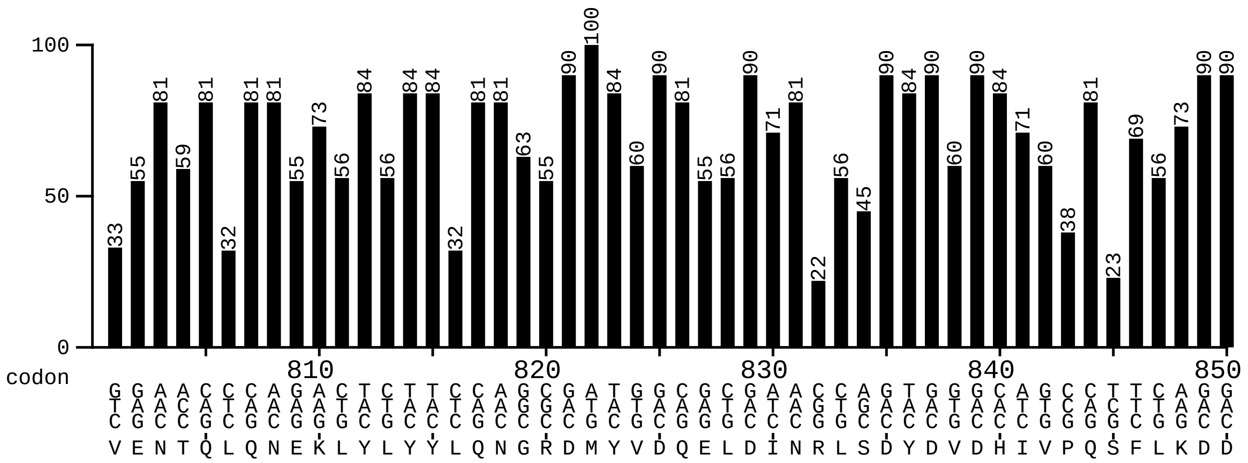
100
50
0
codon
810 820 830 840 850
V E N T Q L Q N E K L Y L Y Y L Q N G R D M Y V D Q E L D I N R L S D Y D V D H I V P Q S F L K D D

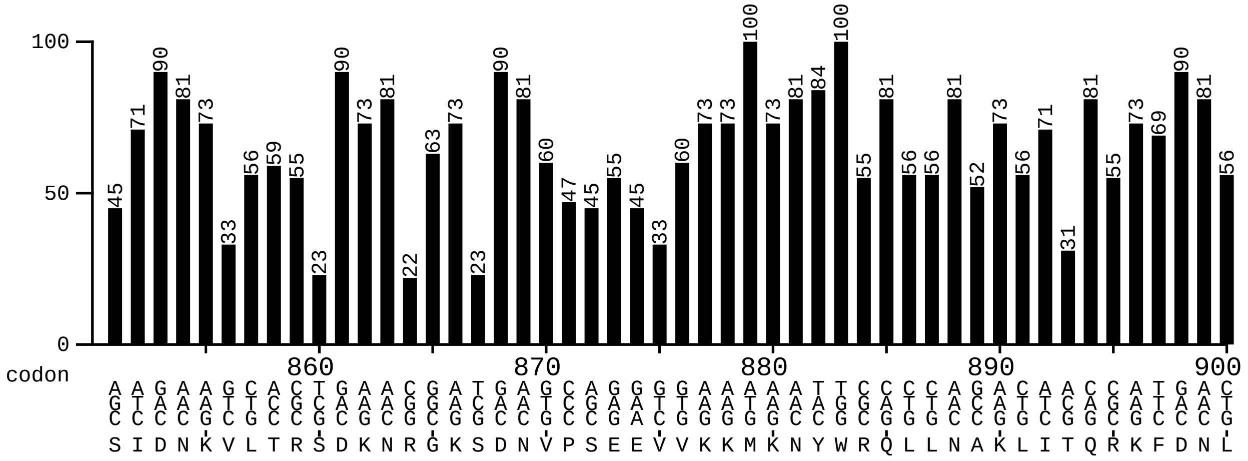
100
50
0
codon
860 870 880 890 900
S I D N K V L T R S D K N R G K S D N V P S E E V V K K M K N Y W R Q L L N A K L I T Q R K F D N L

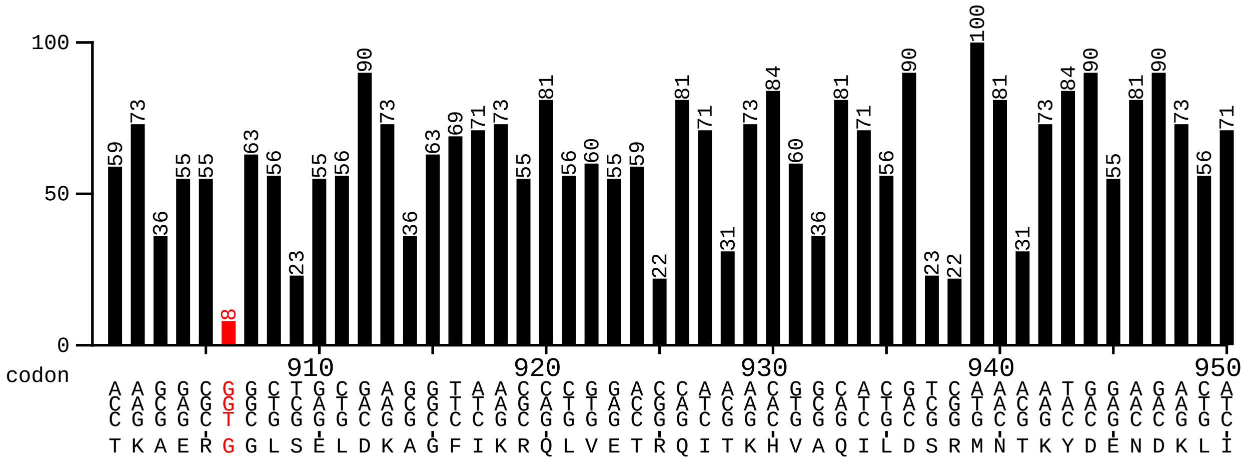
100
50
0
codon
910 920 930 940 950
T K A E R G G L S E L D K A G F I K R Q L V E T R Q I T K H V A Q I L D S R M N T K Y D E N D K L I

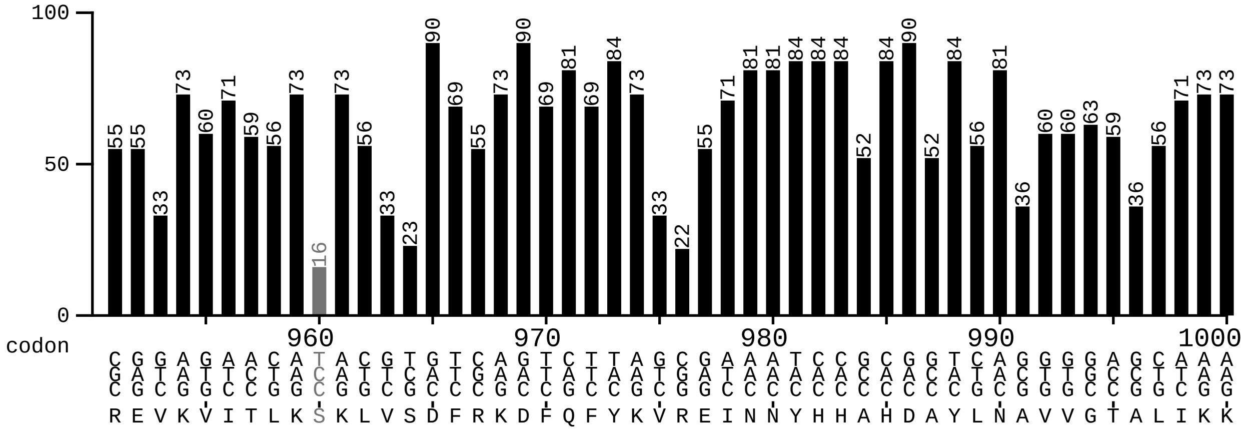

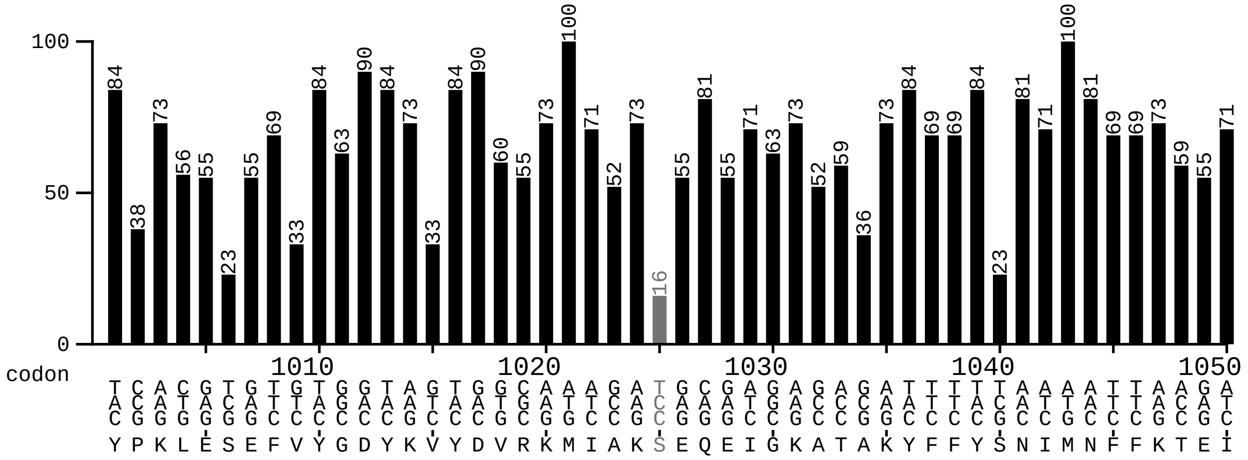

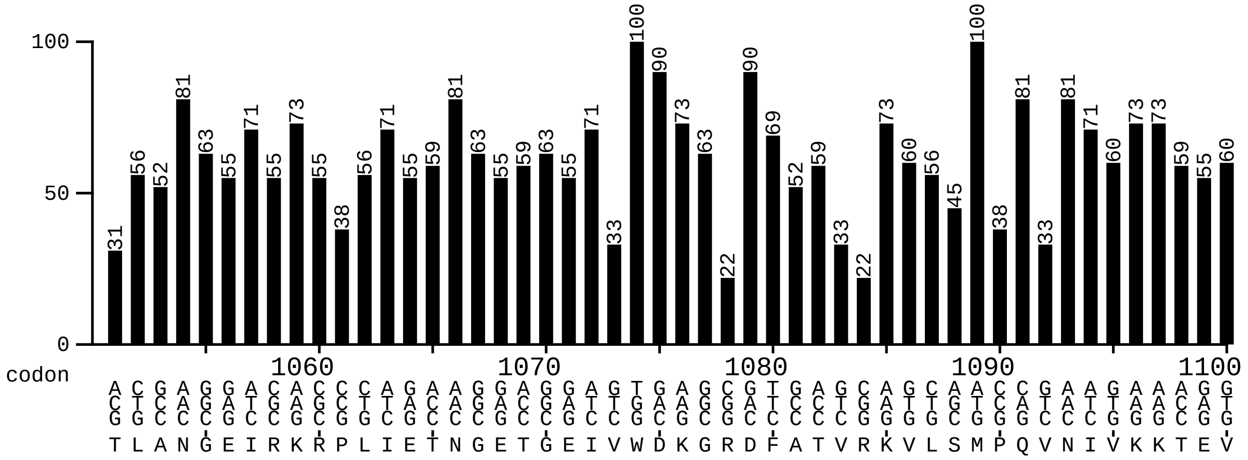

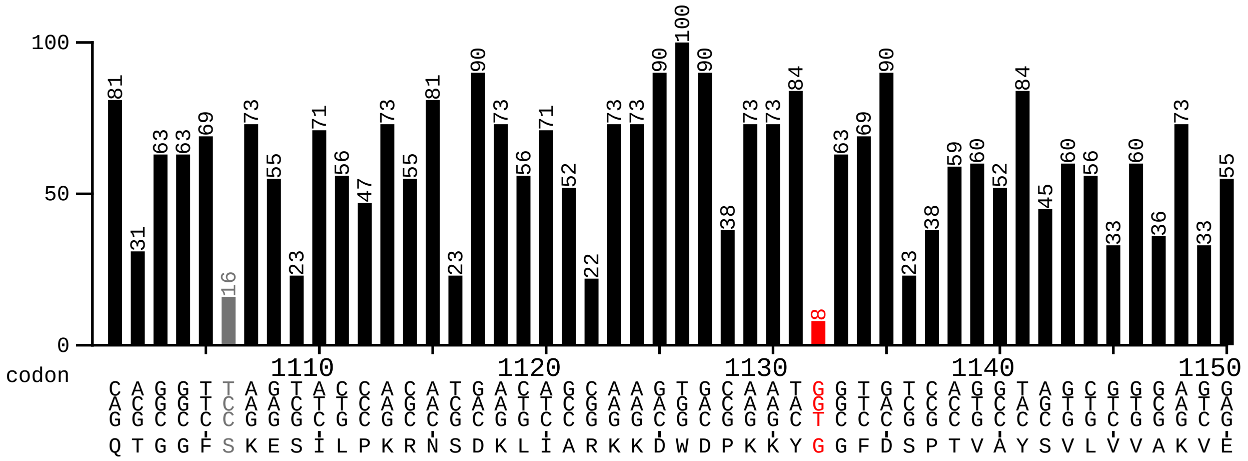

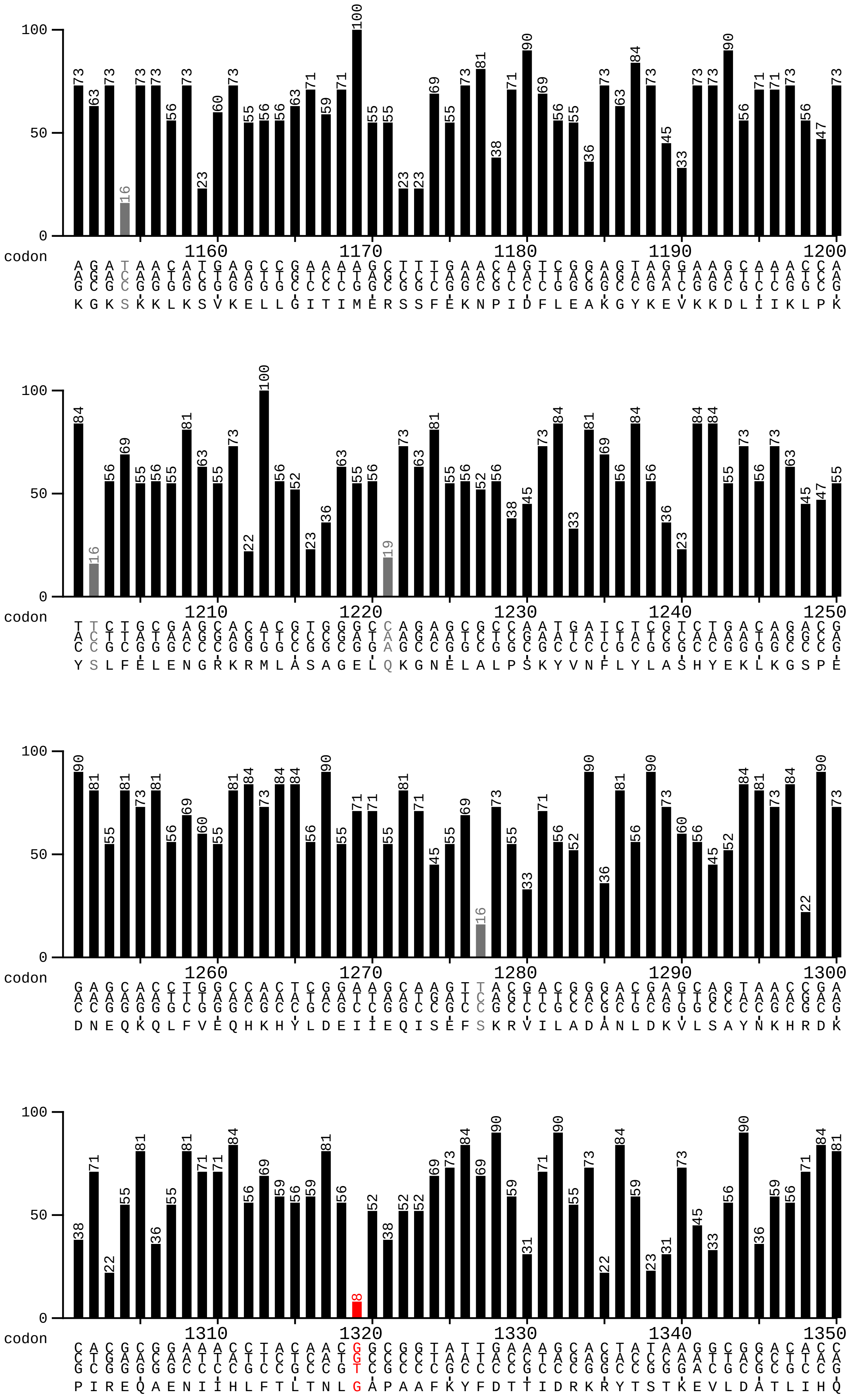

100
50
0
codon
1160 1170 1180 1190 1200
K G K S K K L K S V K E L L G I T I M E R S S F E K N P I D F L E A K G Y K E V K K D L I I K L P K
1210 1220 1230 1240 1250
Y S L F E L E N G R K R M L A S A G E L Q K G N E L A L P S K Y V N F L Y L A S H Y E K L K G S P E
1260 1270 1280 1290 1300
D N E Q K Q L F V E Q H K H Y L D E I I E Q I S E F S K R V I L A D A N L D K V L S A Y N K H R D K
1310 1320 1330 1340 1350
P I R E Q A E N I I H L F T L T N L G A P A A F K Y F D T T I D R K R Y T S T K E V L D A T L I H Q

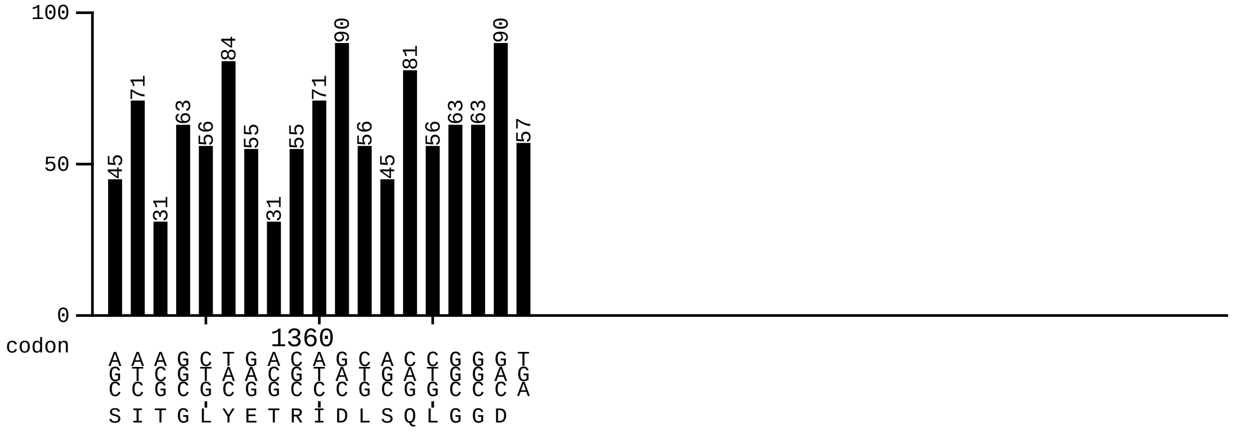
100
50
0
45 71 31 63 56 84 55 31 55 71 90 56 45 81 56 63 63 90 57
codon
1360
AGC ATC ACG GGC CTG TAC GAG ACG CGC ATC GAC CTG AGC CAG CTG GGC GGC GAC TGA
S I T G L Y E T R I D L S Q L G G D

graphical codon usage analyser 2.0

powered by www.mrgene.com
$ 0.39 /bp
0.30 € /bp

www.gcua.de
created: 18.10.2019

Cas9
sequence derived from M. smegmatis

Codontable:
https://www.kazusa.or.jp/codon/cgi-bin/showcodon.cgi?species=190485&aa=1&style=N

Ordinate (y-axis): frequency <20% <10%

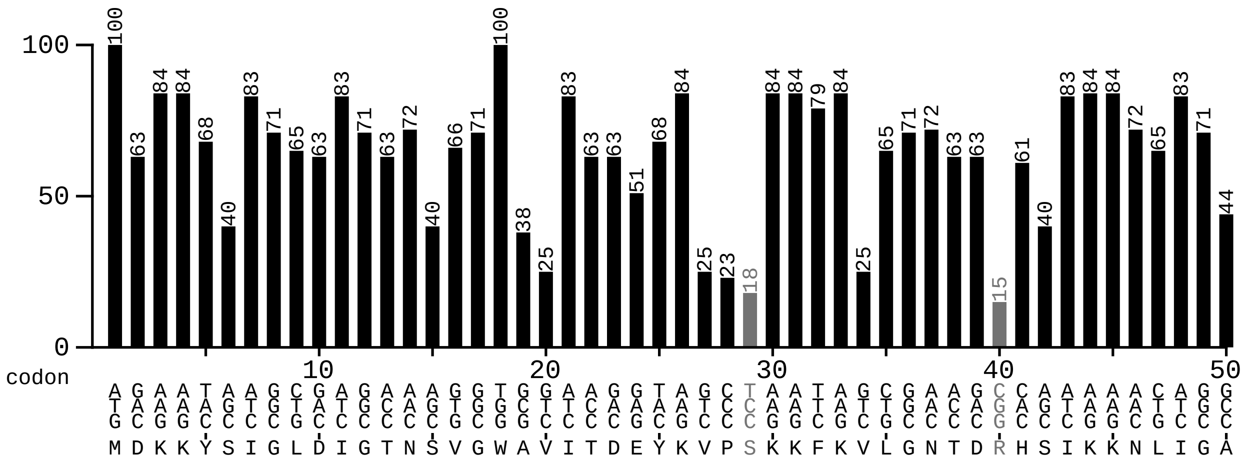


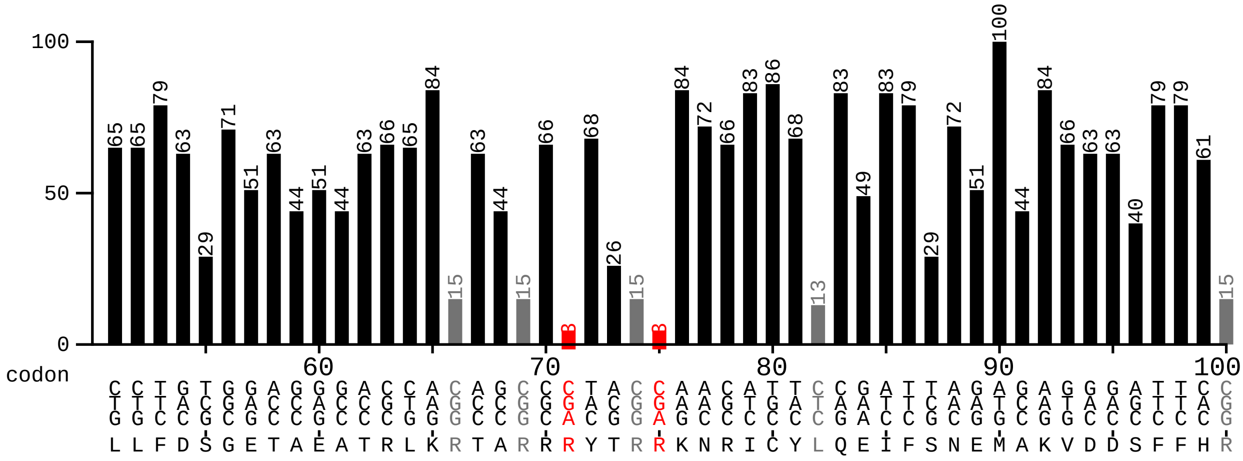


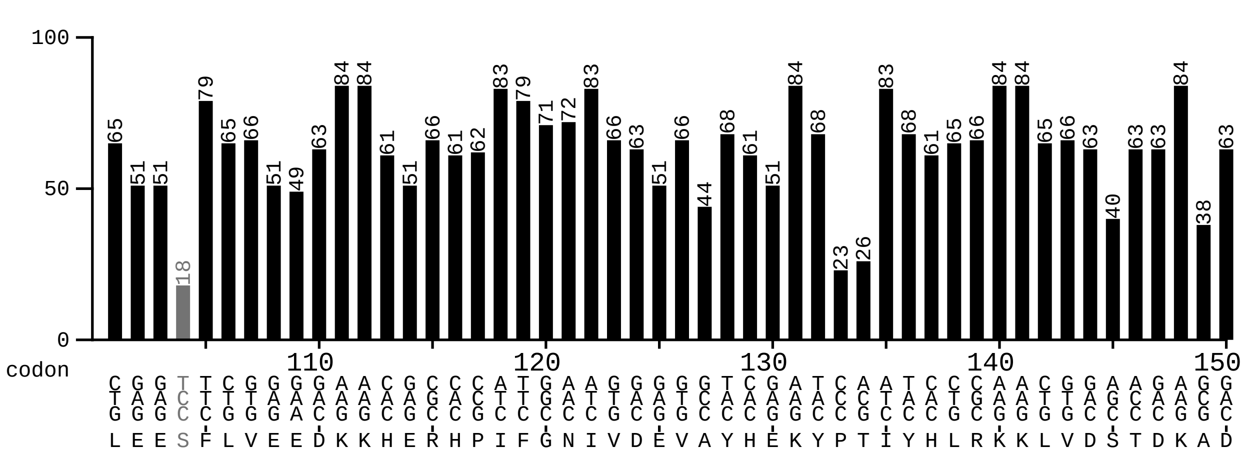

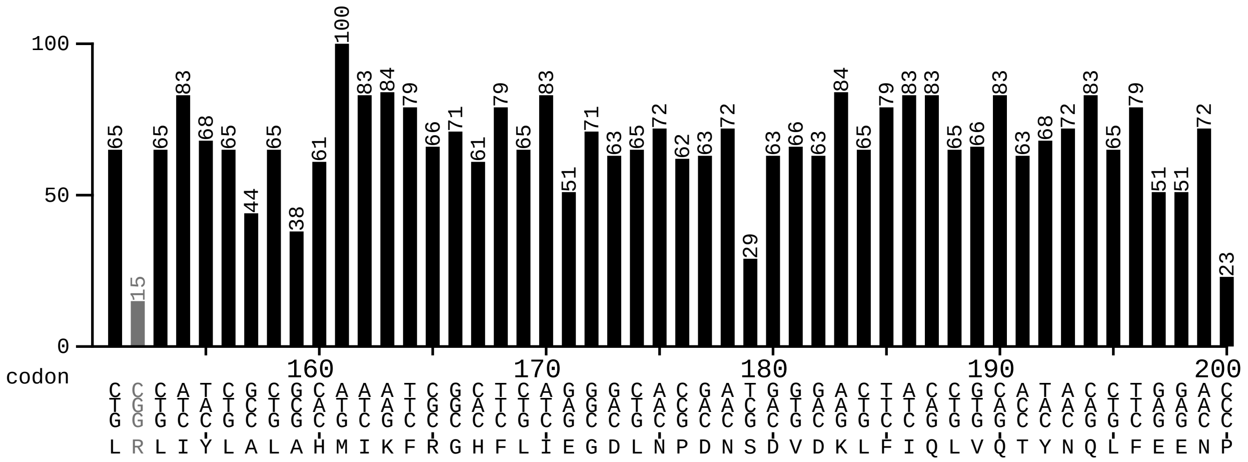
100
50
0
codon
160 170 180 190 200
L R L I Y L A L A H M I K F R G H F L I E G D L N P D N S D V D K L F I Q L V Q T Y N Q L F E E N P

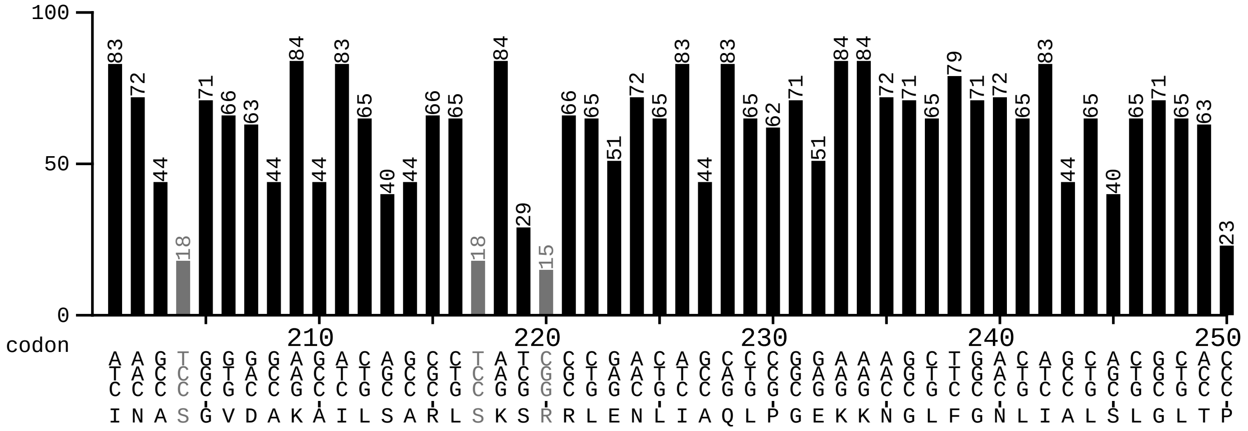
100
50
0
codon
210 220 230 240 250
I N A S G V D A K A I L S A R L S K S R R L E N L I A Q L P G E K K N G L F G N L I A L S L G L T P

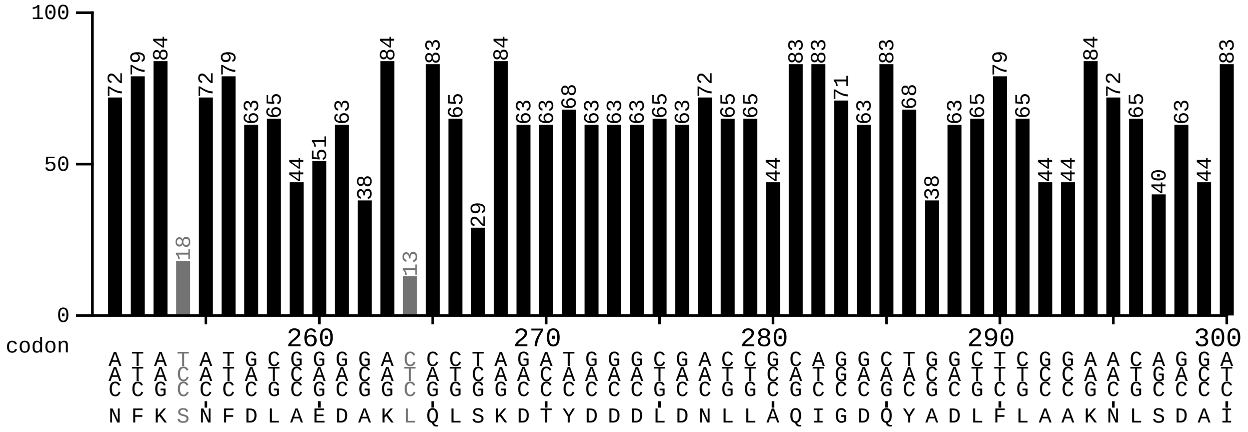
100
50
0
codon
260 270 280 290 300
N F K S N F D L A E D A K L Q L S K D T Y D D D L D N L L A Q I G D Q Y A D L F L A A K N L S D A I

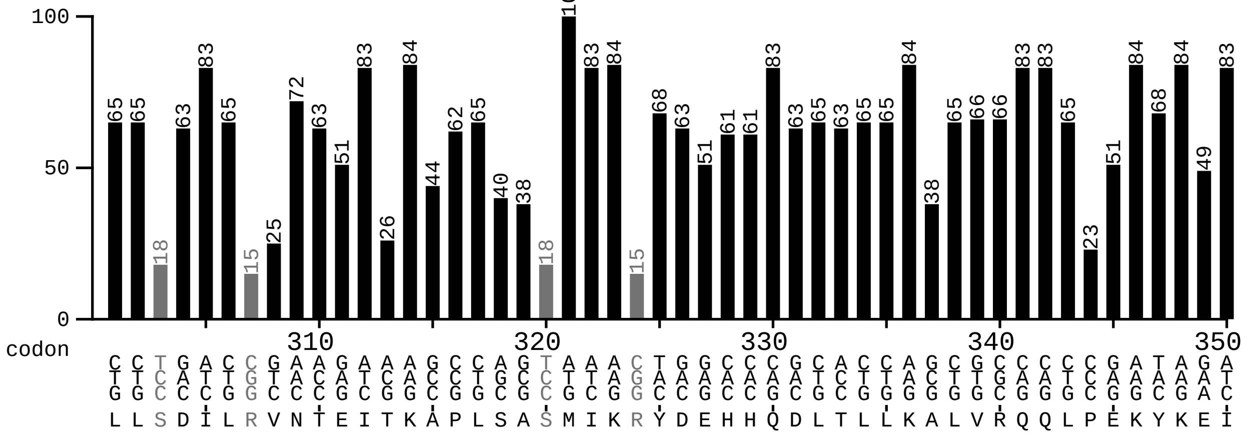
100
50
0
codon
310 320 330 340 350
L L S D I L R V N T E I T K A P L S A S M I K R Y D E H H Q D L T L L K A L V R Q Q L P E K Y K E I

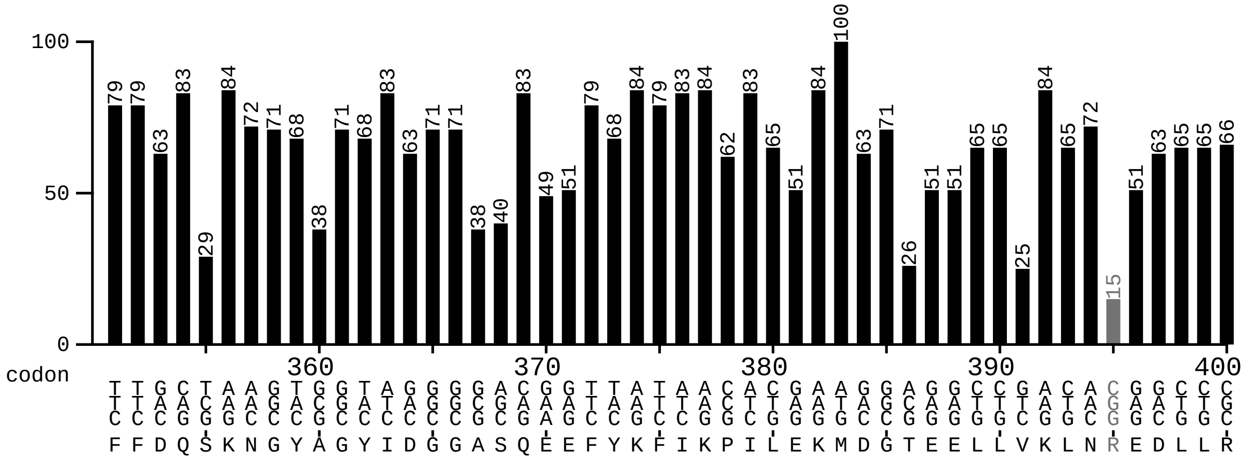
100
50
0
codon
360 370 380 390 400
F F D Q S K N G Y A G Y I D G G A S Q E E F Y K F I K P I L E K M D G T E E L L V K L N R E D L L R

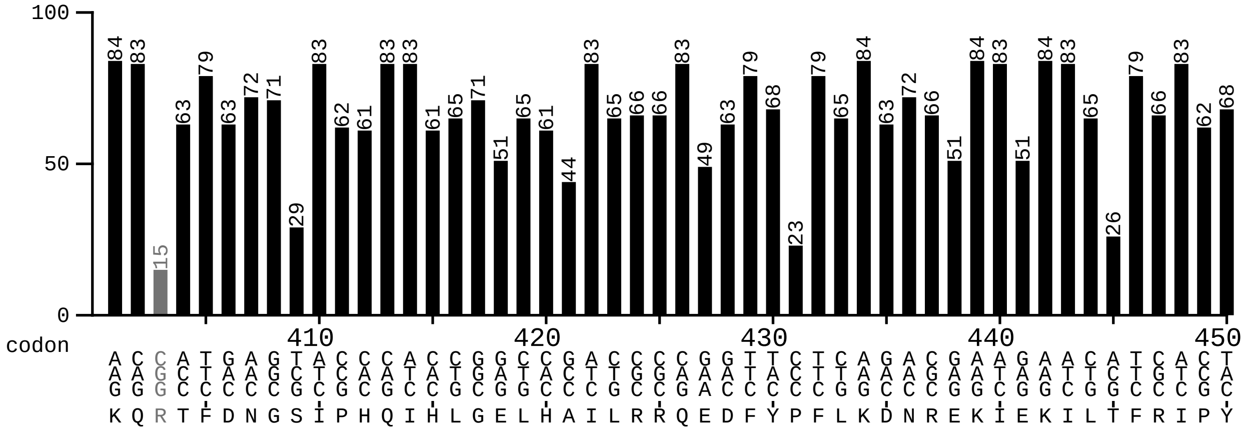
100
50
0
codon
410 420 430 440 450
K Q R T F D N G S I P H Q I H L G E L H A I L R R Q E D F Y P F L K D N R E K I E K I L T F R I P Y

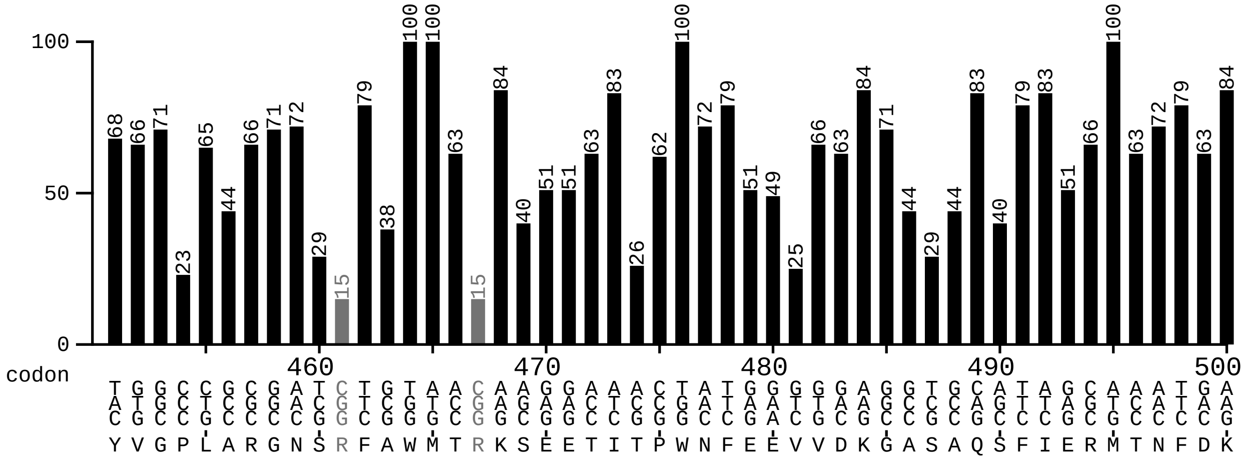
100
50
0
codon
460 470 480 490 500
Y V G P L A R G N S R F A W M T R K S E E T I T P W N F E E V V D K G A S A Q S F I E R M T N F D K

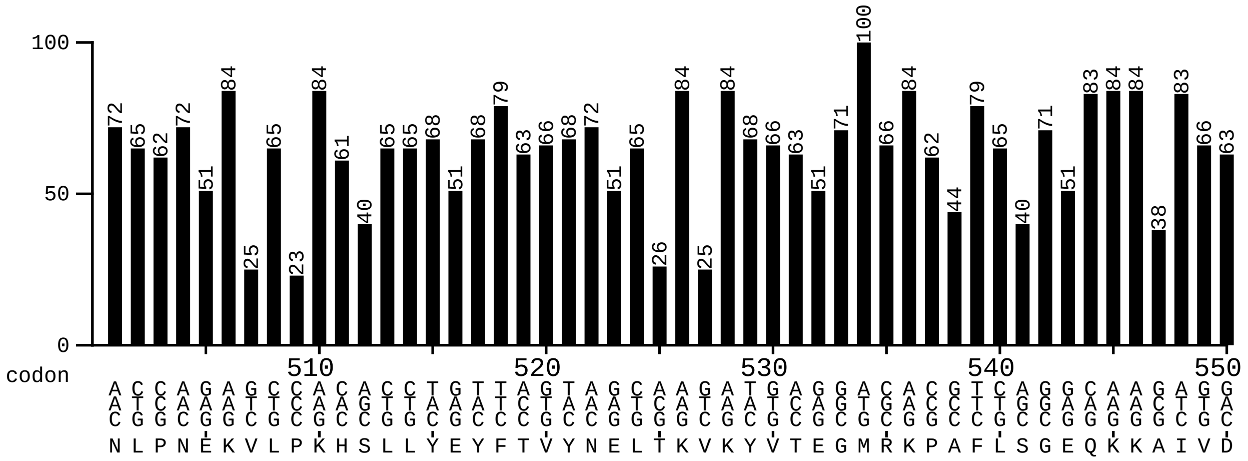
100
50
0
codon
510 520 530 540 550
N L P N E K V L P K H S L L Y E Y F T V Y N E L T K V K Y V T E G M R K P A F L S G E Q K K A I V D

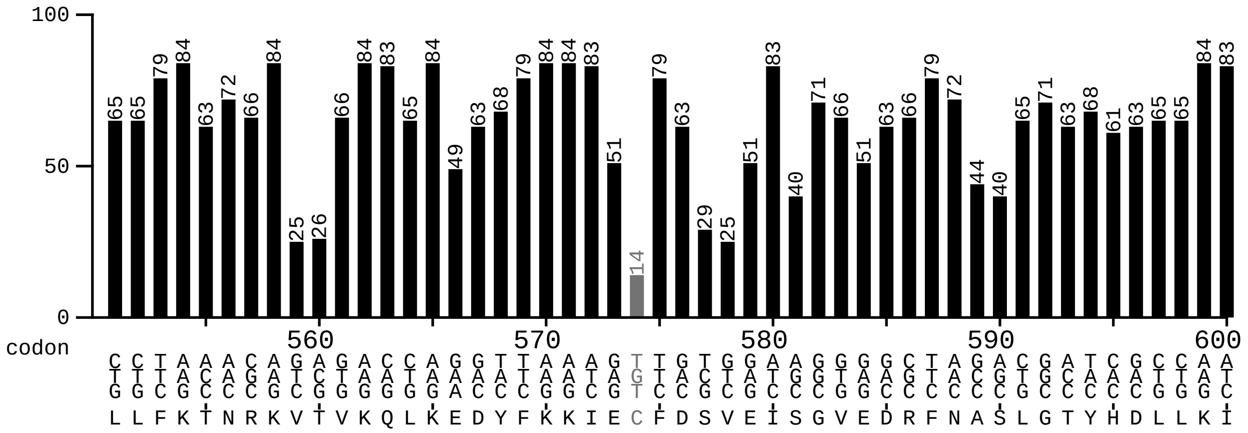
100
50
0
codon
560 570 580 590 600
L L F K T N R K V T V K Q L K E D Y F K K I E C F D S V E I S G V E D R F N A S L G T Y H D L L K I

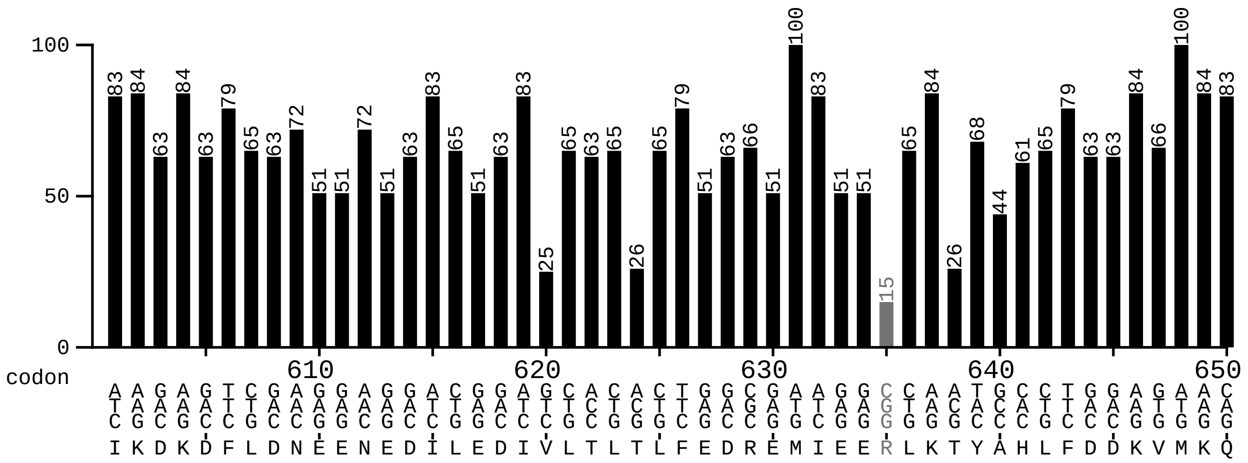
100
50
0
codon
610 620 630 640 650
I K D K D F L D N E E N E D I L E D I V L T L T L F E D R E M I E E R L K T Y A H L F D D K V M K Q

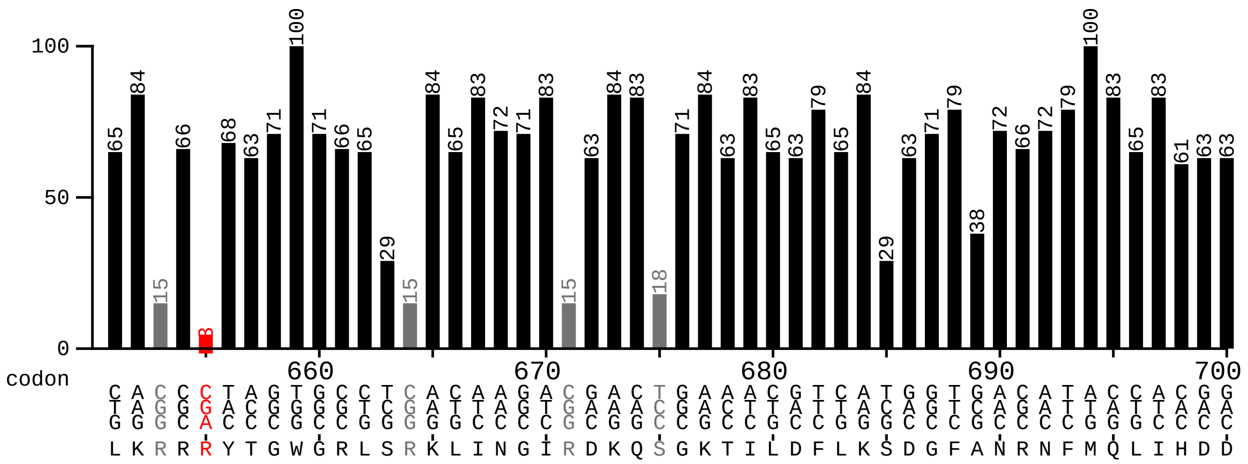
100
50
0
codon
660 670 680 690 700
L K R R R Y T G W G R L S R K L I N G I R D K Q S G K T I L D F L K S D G F A N R N F M Q L I H D D

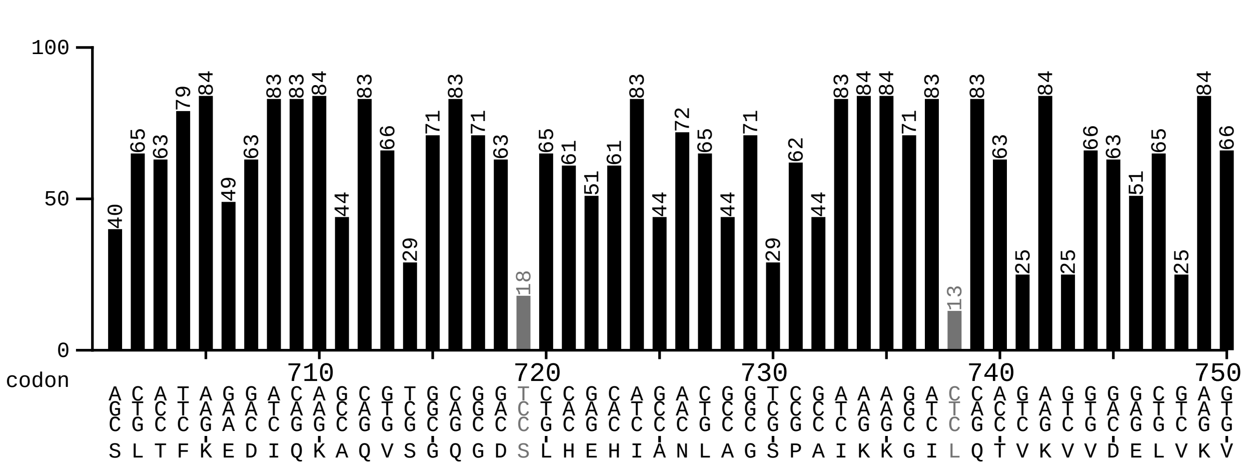
100
50
0
codon
710 720 730 740 750
S L T F K E D I Q K A Q V S G Q G D S L H E H I A N L A G S P A I K K G I L Q T V K V V D E L V K V

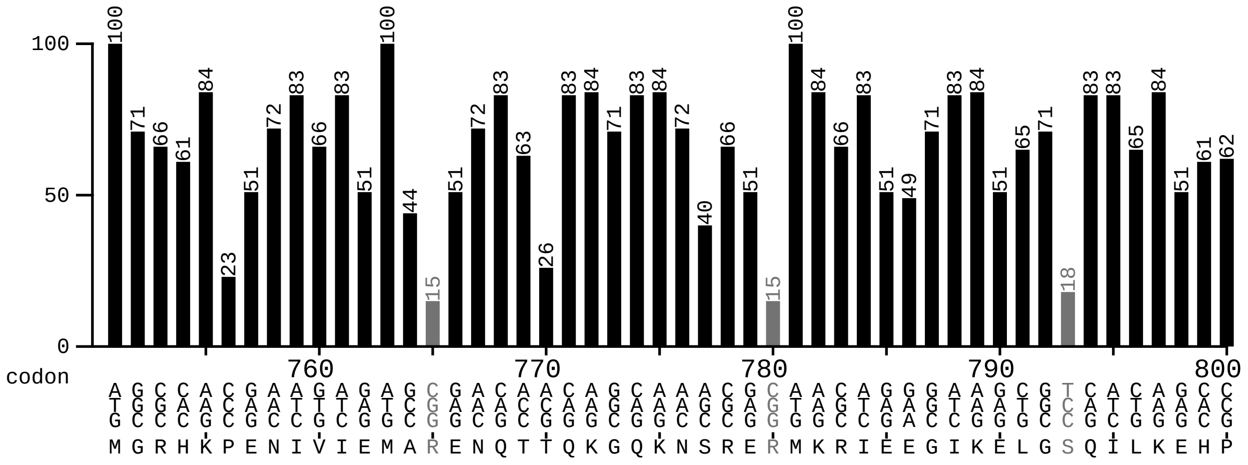
100
50
0
codon
760 770 780 790 800
M G R H K P E N I V I E M A R E N Q T T Q K G Q K N S R E R M K R I E E G I K E L G S Q I L K E H P

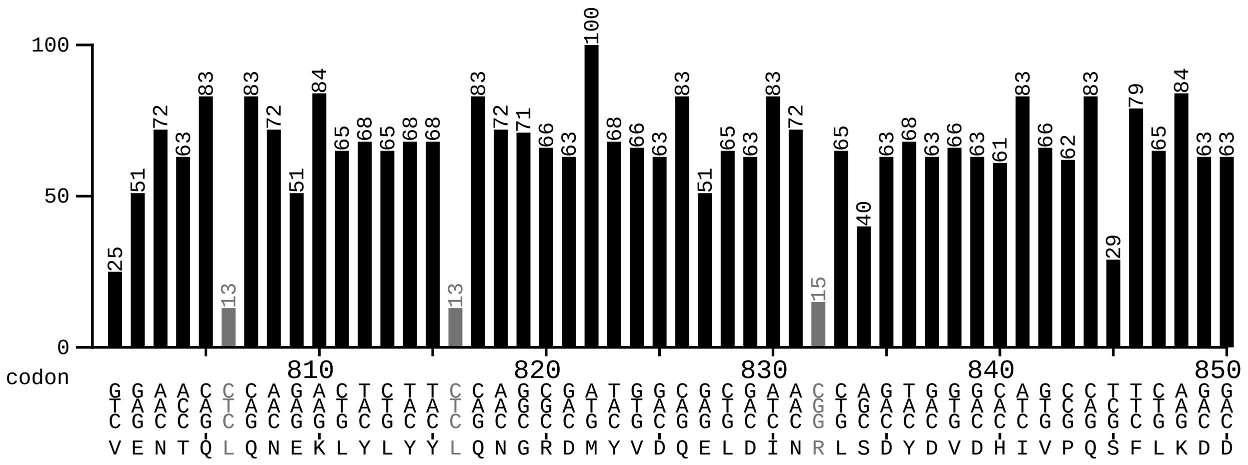
100
50
0
codon
810 820 830 840 850
V E N T Q L Q N E K L Y L Y Y L Q N G R D M Y V D Q E L D I N R L S D Y D V D H I V P Q S F L K D D

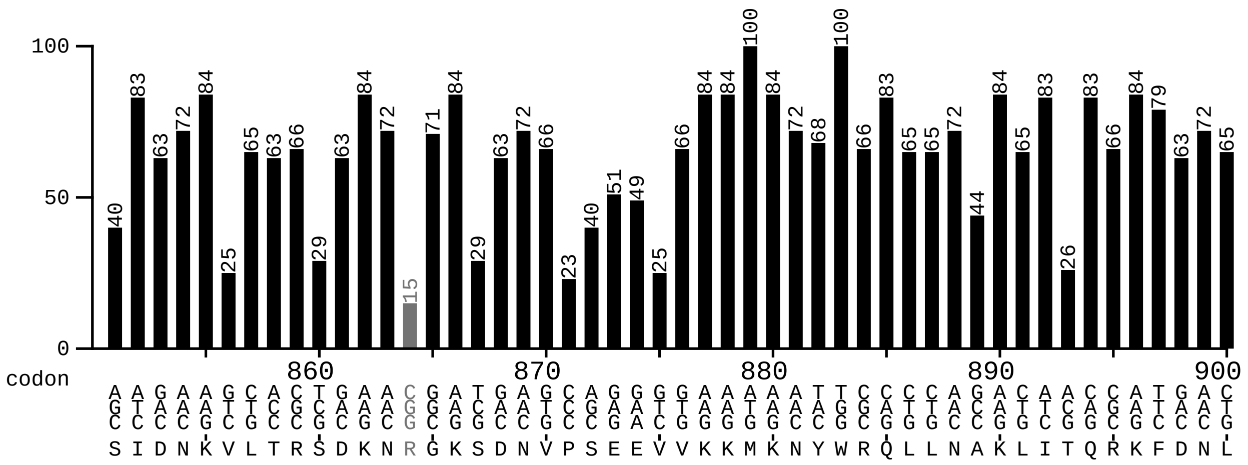
100
50
0
codon
860 870 880 890 900
S I D N K V L T R S D K N R G K S D N V P S E E V V K K M K N Y W R Q L L N A K L I T Q R K F D N L

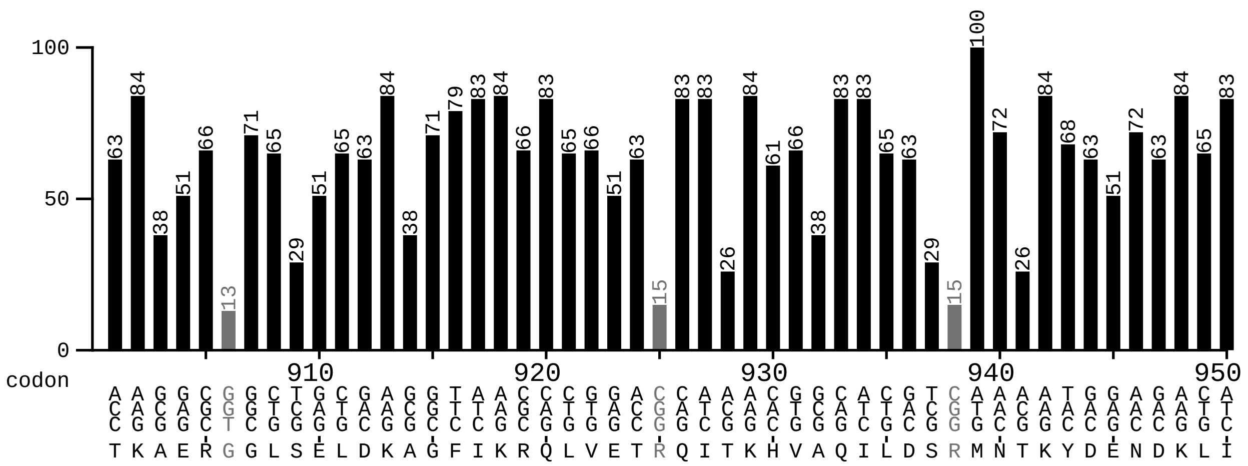
100
50
0
codon
910 920 930 940 950
T K A E R G G L S E L D K A G F I K R Q L V E T R Q I T K H V A Q I L D S R M N T K Y D E N D K L I

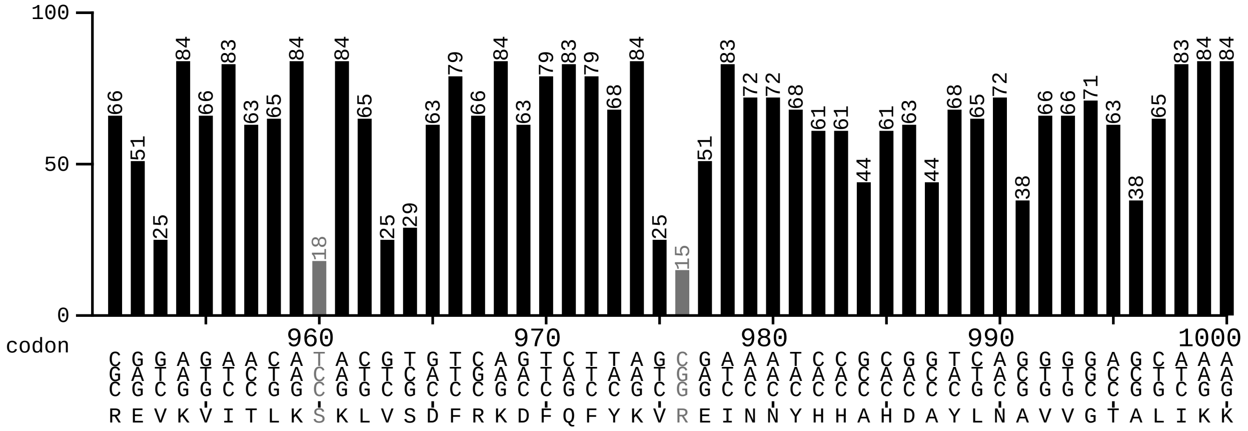
100
50
0
codon
960
970
980
990
1000

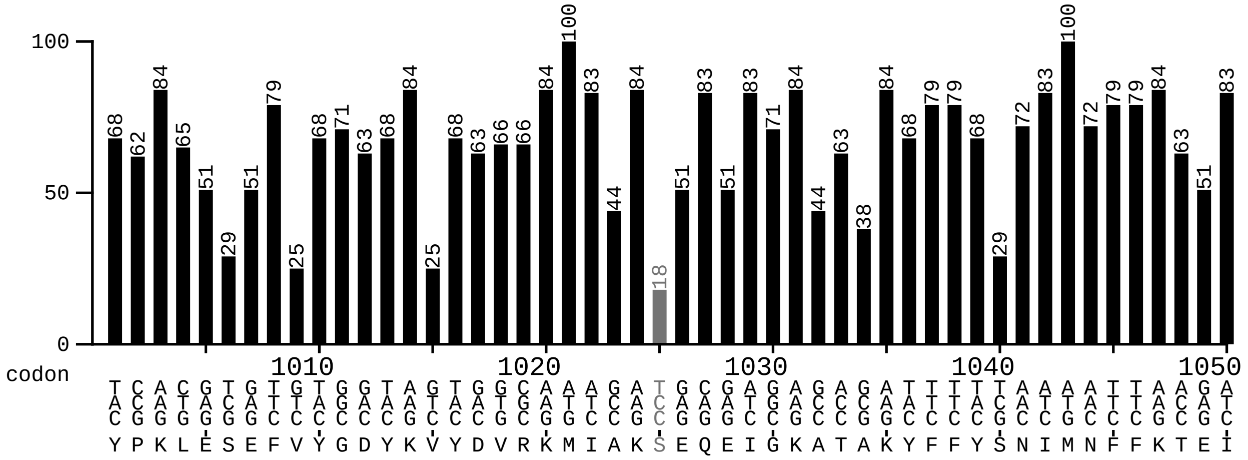
100
50
0
codon
1010
1020
1030
1040
1050

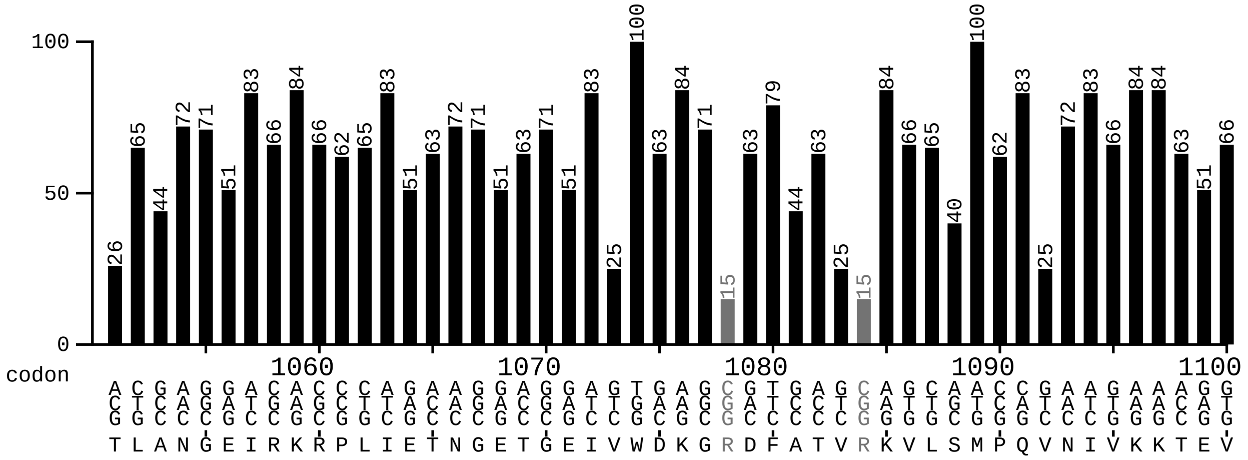
100
50
0
codon
1060
1070
1080
1090
1100

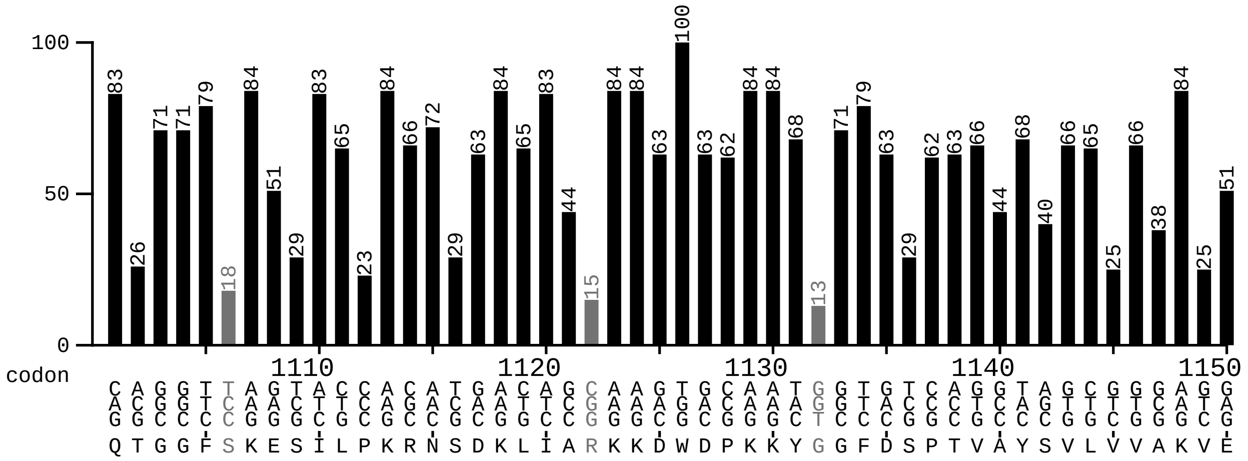
100
50
0
codon
1110
1120
1130
1140
1150

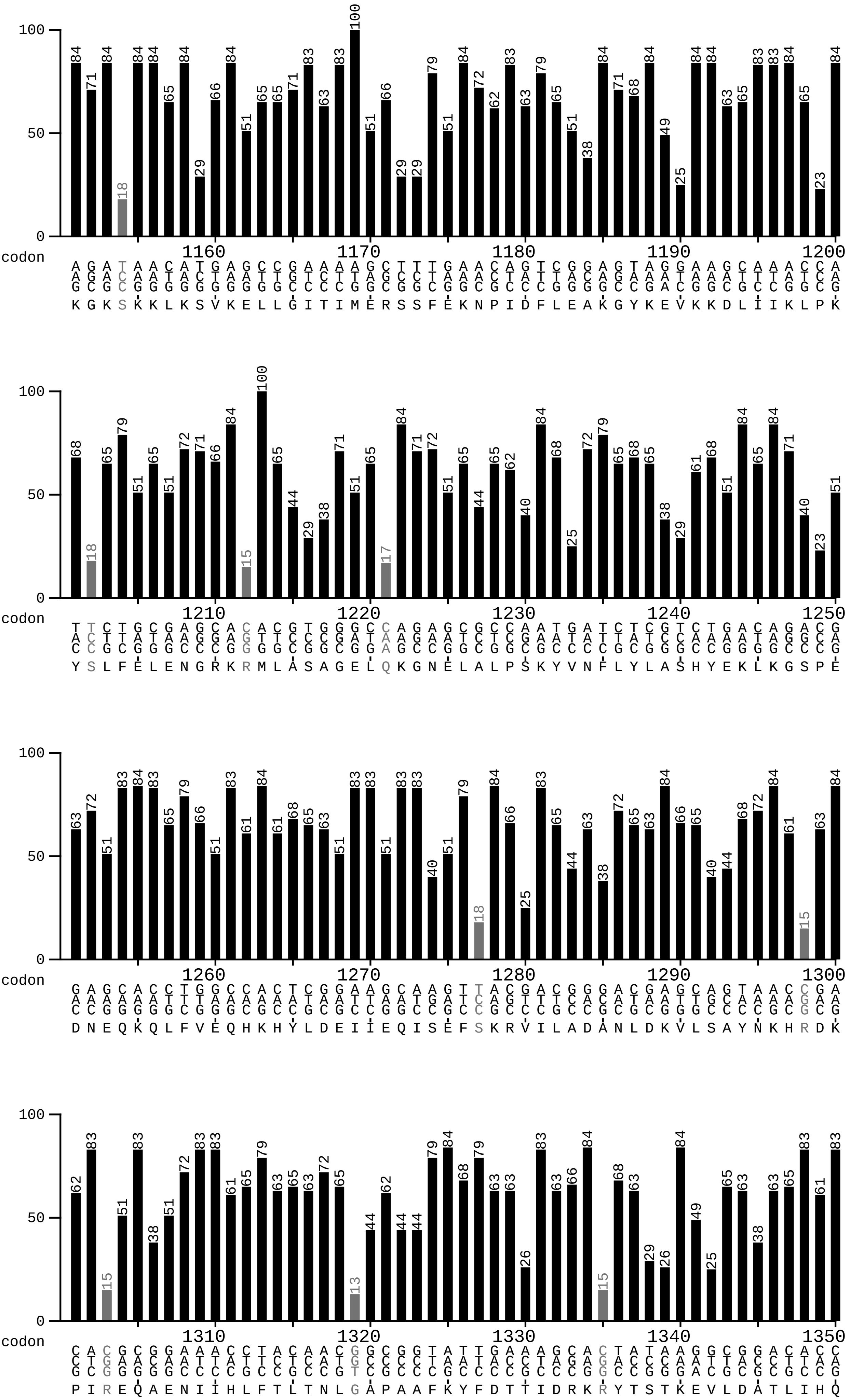

100
50
0
codon
1160
1170
1180
1190
1200
1210
1220
1230
1240
1250
1260
1270
1280
1290
1300
1310
1320
1330
1340
1350

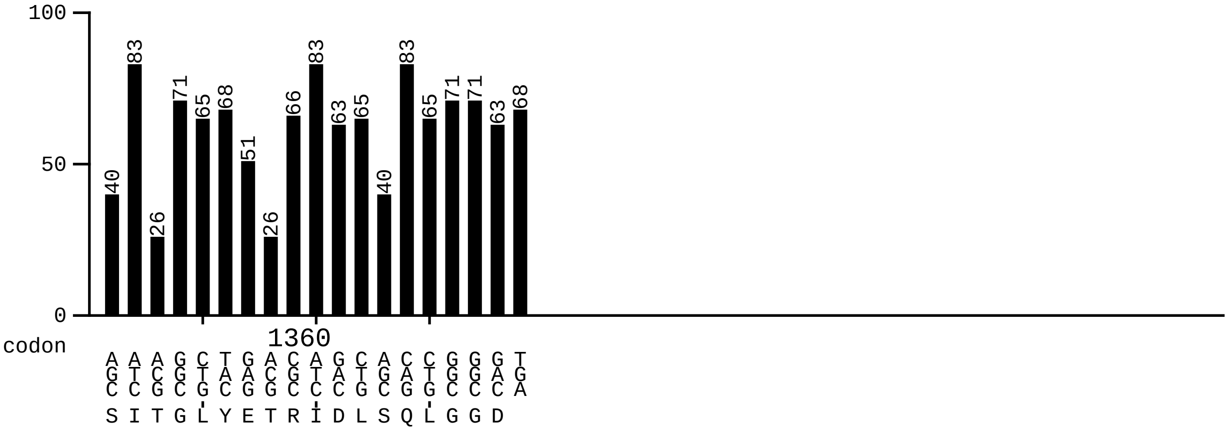
100
50
0
codon
1360
40 83 26 71 65 68 51 26 66 83 63 65 40 83 65 71 71 63 68
AGC ATC ACG GGC CTG TAC GAG ACG CGC ATC GAC CTG AGC CAG CTG GGC GGC GAC TGA
S I T G L Y E T R I D L S Q L G G D
